# Global epistasis and the emergence of ecological function

**DOI:** 10.1101/2022.06.21.496987

**Authors:** Juan Diaz-Colunga, Abigail Skwara, Jean C. C. Vila, Djordje Bajic, Álvaro Sánchez

## Abstract

The emergence of community functions is the result of a complex web of interactions between organisms and their environment. This complexity poses a significant obstacle in quantitatively predicting ecological function from the species-level composition of a community. In this study, we demonstrate that the collective impact of interspecies interactions leads to the emergence of simple linear models that predict ecological function. These predictive models mirror the patterns of *global epistasis* reported in genetics, and they can be quantitatively interpreted in terms of pairwise ecological interactions between species. Our results illuminate an unexplored path to quantitatively linking the composition and function of ecological communities, bringing the tasks of predicting biological function at the genetic, organismal, and ecological scales under the same quantitative formalism.

---

Ecological communities carry out critical functions in both natural and biotechnological settings, from nutrient cycling in the soils (*1*) to biofuel production in industrial biorefineries (*2*). Manipulating community-level functions is therefore a major aspiration across multiple research fields and sectors of the economy. This aspiration is currently hampered by the lack of quantitative models that predictively link the species-level composition of a community with the functions it provides. Lacking such general predictive models we cannot determine, for instance, which particular phages in a cocktail optimize its antibacterial activity (*3*), or which combination of plant species maximizes the harvested yield in a mixed crop setting (*4*) (Fig. 1A). Answering these questions requires us to quantitatively connect the composition and the function of ecological communities. This is a formidable challenge: community-level functions generally emerge from a complex web of molecular, physiological, and organismal interactions, which can be difficult to characterize and vary substantially across ecological contexts. Because of this complexity, the effect of a particular species on the function of a community will generally depend on which other species are present, making it difficult to predict.

**Fig. 1.**
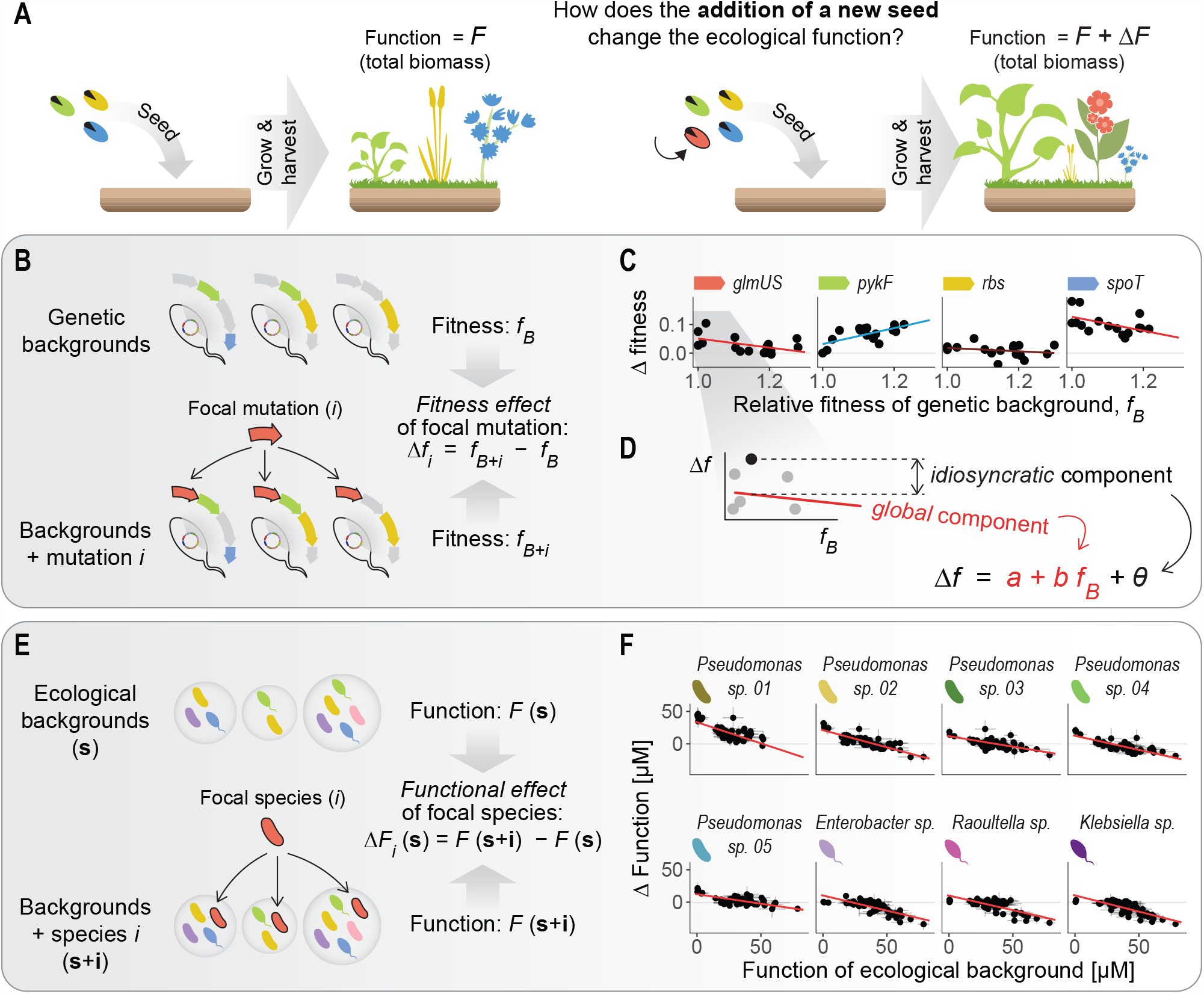
An ecological parallel to global epistasis. (**A**) The central challenge of this work is predicting how a collective ecological function will change with the addition of a species. (**B**-**C**) Recent work in quantitative genetics has found that the fitness effect of a mutation is often predicted by the fitness of the genetic background where it arises, following simple linear regressions. This phenomenon has been termed *global epistasis*. The slope and intercept of the linear regression vary across different mutations. Data from Khan et al. (*7*) (**D**) The fitness effect of a mutation can thus be broken down into two components: first, a *global* component that is predictable from the linear model, and, second, an *idiosyncratic* component that is not predictable from the background fitness, represented by the residuals of the fits. (**E**) We hypothesized the existence of an ecological parallel to global epistasis, where the addition of a focal species to a community might induce a change in a community-level function that can be predicted from the function of the background community where the focal species is added. (**F**) We inoculated different combinations of eight bacterial isolates in synthetic liquid medium. The functions of these consortia were quantified as the level of pyoverdines secretion after 48 h of incubation (Materials and Methods, fig. S1). We found that the *functional effect* of adding a species to an *ecological background* was well predicted by a linear model linking it to the function of the background community.

The challenges of connecting biological structure to function are not exclusive to ecology. In genetics, for instance, the contribution of a locus to a quantitative phenotype depends on its genetic background, via genetic (*epistatic*) interactions with other loci (*5*). Encouragingly, while these interactions can also be highly complex, recent research has found that the fitness effect of a mutation is often well predicted by the fitness of its genetic background through simple, linear quantitative models (Fig. 1B-D). The emergence of these simple linear patterns is a manifestation of *global epistasis* (*6*–*17*) — a phenomenon that includes (but is not limited to) the common observation that beneficial mutations have smaller fitness effects in higher-fitness genetic backgrounds (*diminishing returns epistasis*). As an illustration of global epistasis, in Fig. 1C we reproduce previous results showing that the fitness effects of four *E. coli* mutations are well predicted by the fitness of their genetic backgrounds through simple linear regressions (*7*). These global epistasis patterns can be determined from just a small number of empirical observations, and theory has shown that they can be mechanistically interpreted in terms of microscopic gene-by-gene interactions (*16,17*). Recent studies have leveraged global epistasis to develop highly promising methodologies for inferring full genotype-phenotype maps in large combinatorial spaces from just a subset of measurements (*18*–*22*).

Inspired by recently established parallels between genetic and functional ecological interactions (*23*–*25*), here we hypothesize that an ecological analog to global epistasis might exist: simple linear regressions might predict how adding a species to a community will affect its community-level function (Fig. 1E). If such regressions existed and could be interpreted in terms of interspecies interactions (as is the case for global epistasis in genetics), this would unlock our ability to predictively connect species-level composition to quantitative function across a variety of ecological systems.

To evaluate the merits of this hypothesis, we conducted a series of experiments using synthetic microbial communities and also examined previously published data of plant, bacterial, and algal ecosystems, under distinct environmental conditions and for a variety of collective functions. We found that a parallel concept to global epistasis can indeed be formulated in all these cases. Here we show that this allows us to build predictive models to optimize ecological function which are simple, general, and highly interpretable. Furthermore, we extend previous theoretical results from the field of genetics to demonstrate that ecological global epistasis-like patterns can be quantitatively linked to microscopic species-by-species interactions. Our findings argue that the same general formalism can be applied to predict biological function across widely different scales of organization, from molecules and organisms to ecological communities.

## An ecological parallel to global epistasis predicts the functional effect of a species

Based on our hypothesis, we could define the *functional effect* of a species by analogy with the fitness effect of a mutation, i.e., as the difference in function between two communities that differ solely in the presence or absence of such species (Fig. 1E). If our hypothesis were correct, we should observe that the functional effect of a focal species would be well predicted by the function of the (background) community where we include it, following simple linear models similar to those observed in genetic global epistasis.

To test our hypothesis, we built a small library of eight soil bacterial isolates (Materials and Methods), from which one could potentially assemble 255 different consortia based on the presence/absence of each member. The community-level function we studied was the net production of pyoverdines. This represents a good test case for an ecological function that is sensitive to species interactions, as pyoverdine secretion is known to respond to intra-species signaling (*26*) and it is also often controlled by population size via quorum sensing (*27*). Five of our species were *Pseudomonas* strains that produce pyoverdines in monoculture, while the remaining three were non-producing *Enterobacteriaceae* (fig. S1).

From this library, we assembled a subset (*N* = 164) of all possible unique species combinations, including an approximately even representation of pairs, trios, etc. (Supplementary Text). Each of these consortia can be represented by a vector **s**, which encodes the presence/absence of each species *i* (*s*_*i*_ = 0, 1). To form each consortium, we inoculated every member at a fixed inoculum density in minimal liquid growth medium (fig. S1, Materials and Methods). We then incubated our consortia for 48 h and measured their function (*F* (**s**)) as the concentration of pyoverdines in the spent media (fig. S1, Materials and Methods). The assembled consortia exhibited high variation in functional levels, with pyoverdine concentrations ranging from 0 to 70 µM (fig. S1). In Fig. 1F we plot the functional effect of each species (Δ*F*_*i*_ (**s**) = *F* (**s** + **i**) − *F* (**s**) for species *i*, Fig. 1E) against the function of its background consortia (*F* (**s**)).

Consistent with our hypothesis, we found that the functional effects of all species in different community contexts were well predicted by simple linear relationships of the form Δ*F*_*i*_ (**s**) = *a*_*i*_ + *b*_*i*_ *F* (**s**) + *θ*_*i*_ (**s**) (Fig. 1F). We call this expression the *functional effect equation* (FEE) of species *i*. The intercepts (*a*_*i*_) and slopes (*b*_*i*_) of the FEEs differ across taxa, suggesting that they are determined by specific interactions between each individual species and the rest of its ecological partners (fig. S2). The terms *θ*_*i*_ (**s**) (i.e., the residuals of the fits) capture the component of said interactions that is not predictable from the linear fit itself. Consistent with our initial hypothesis, these results indicate that a parallel to global epistasis exists in this ecological system, which predicts how including a particular species into different community contexts will affect their community-level function.

## The ecological parallel to global epistasis is ubiquitous across a wide range of ecological communities

To determine how general this phenomenon might be beyond our particular experimental setting, we re-analyzed a collection of already published datasets where the quantitative relationship between community composition and function had been measured in a similar manner. Table S1 summarizes the datasets we considered, all of which include multiple combinatorial assemblages of species from candidate pools of 4 to 25 taxa. These datasets include both plant (*28*) and phytoplankton (*29*) communities, as well as synthetic bacterial communities formed by either Gram-negative or Gram-positive bacteria (*25,30,31*). This set of communities were assembled under widely different ecological conditions, including the number of organismal generations, the type and frequency of resource addition, and the form of propagation. The functions were also different in each case, ranging from the production of biomass to the net metabolic activity, and from the secretion of specific enzymes to the degradation of environmental polymers. As shown in Fig. 2, simple linear models predict the functional effect of species across all datasets investigated, explaining on average 75% of the functional variance (fig. S3).

**Fig. 2.**
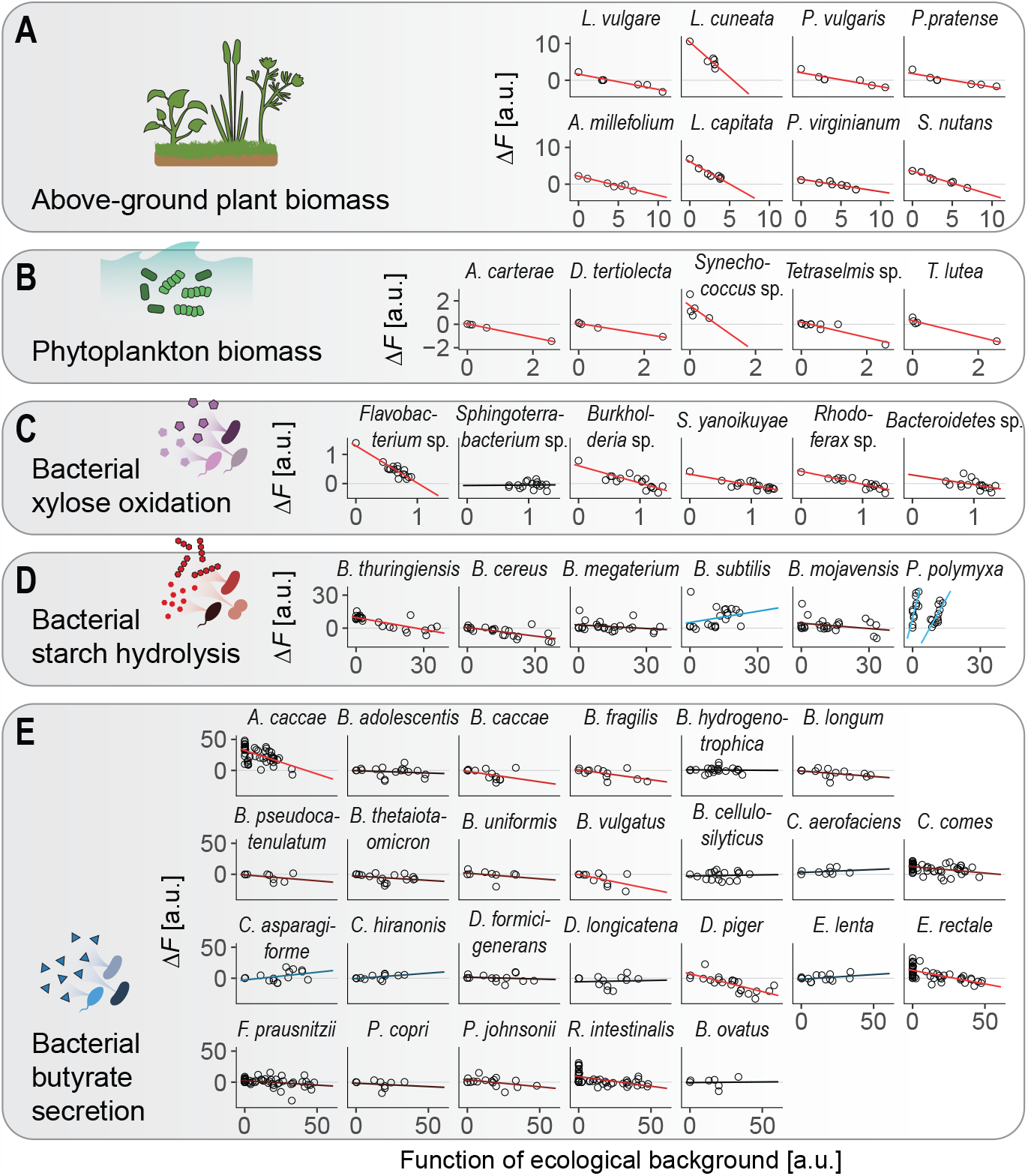
Ecological global epistasis across different communities and functions. The functional effect of a focal species can often be predicted from a simple, linear model linking it to the function of the background community where it is included (red lines: negative slopes, blue lines: positive slopes, black lines: flat slopes). This is observed across communities formed by very different organism types, under different ecological conditions, and for different collective functions (table S1). (**A**) Data from Kuebbing et al. (*28*) (**B**) Data from Ghedini et al. (*29*) (**C**) Data from Langenheder et al. (*30*) (**D**) Data from Sanchez-Gorostiaga et al. (*25*) (**E**) Data from Clark et al. (*31*)

Most species (∼50% across all datasets) display negatively sloped FEEs (red lines in Fig. 2). This trend is also commonly observed in population genetics: the fitness effect of a mutation most often becomes either less beneficial (*diminishing returns*)or more deleterious (*increasing costs*) as the fitness of the genetic background increases (*6–8,11,13,14*). Often, species increase community function when they are included in low performing ecological backgrounds but decrease it when the background function is high. About 45% of all species exhibit relatively flat slopes (black lines in Fig. 2). Note, however, that these flat patterns are also informative for predictive purposes. The magnitude of the deviations from the FEE (even if flat) are useful to discern between (a) species whose contribution to ecosystem function is additive and largely independent of their ecological background (i.e., those species for which the residuals are small), or (b) species whose contribution to the community function depends on the specific composition of their ecological background in a very idiosyncratic manner (i.e., those with large residuals). Finally, a smaller number of species (∼5%) exhibit positively sloped FEEs (blue lines in Fig. 2), becoming more beneficial (or less deleterious) in backgrounds with higher functions. We refer to these patterns as *increasing returns* (or *decreasing costs*).

## Emergence of global functional interactions from pairwise interactions between species

Our analyses suggest that ecological global epistasis is ubiquitous across a wide range of communities. How should we explain this mechanistically? And what factors determine whether a species will exhibit diminishing or accelerating returns? In quantitative genetics, global epistasis patterns may emerge from pairwise epistatic interactions: The slope of the linear regression model for a focal mutation can be approximated by a sum of its effective pairwise interactions with all other mutations in the genetic background (*16,17*) (Supplementary Text). Mutations that engage primarily in negative pairwise epistasis will tend to exhibit diminishing returns, whereas those that engage in positive pairwise epistasis will exhibit increasing returns (*17*). We hypothesized that a similar relationship might hold in ecology, connecting the FEE of a species to its pairwise functional interactions. An effective functional interaction (which we denote as 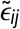 for species *i* and *j*) would capture how, on average, the functional contribution of a pair of species deviates from the sum of the additive effects of each (Fig. 3B, Materials and Methods, Supplementary Text). Extending the theory of global epistasis to community function, the slope (*b*_*i*_) of a species’ FEE could thus be written as (Fig. 3A-C):

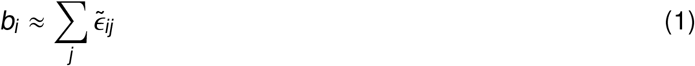

**Fig. 3.**
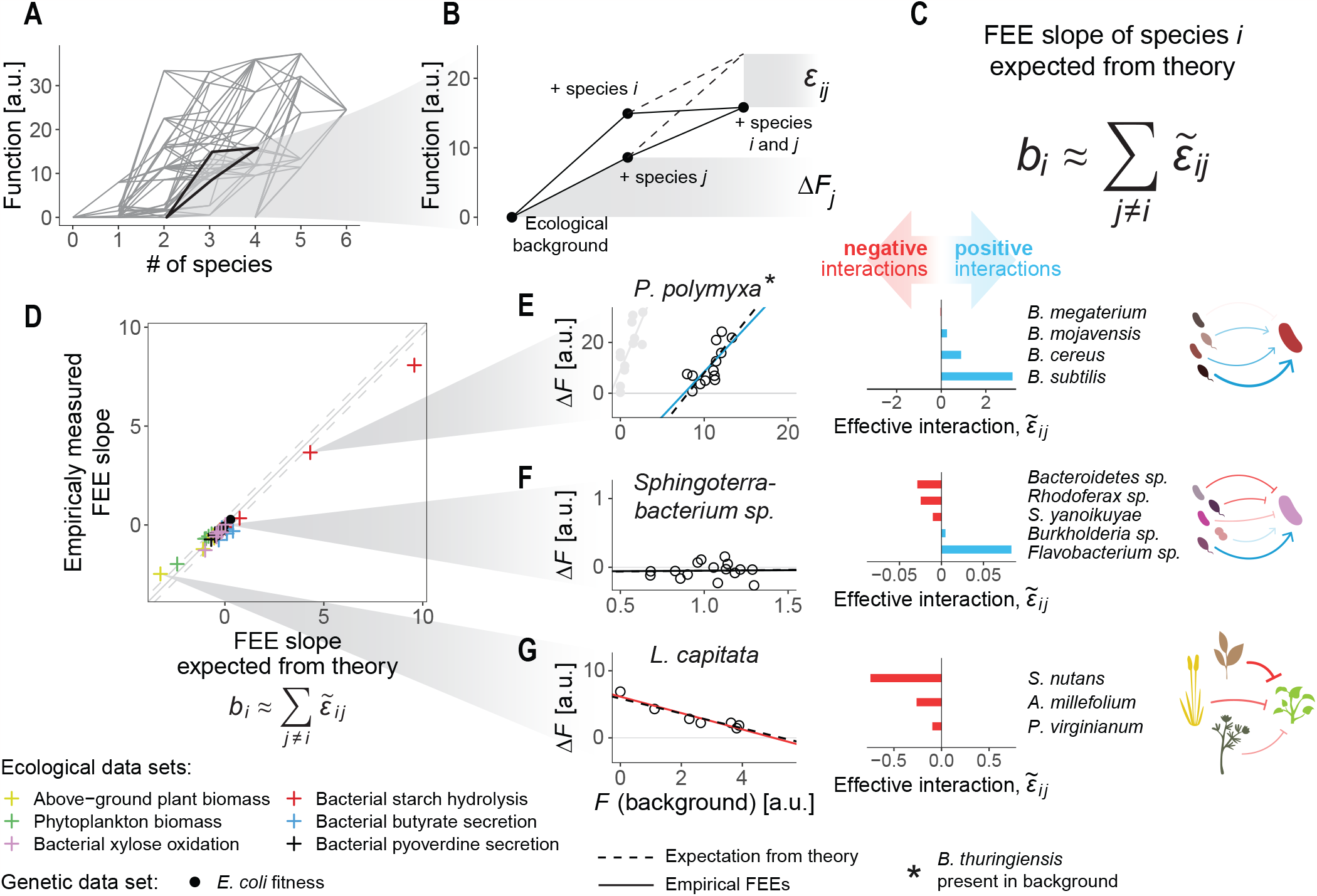
Effective ecological interactions explain FEE slopes. (**A**) Mapping between the number of species and the functions of the communities in the Sanchez-Gorostiaga et al. experiment (*25*). Each node represents a consortium, and edges connect consortia that differ in the presence of a single particular taxon. (**B**) Detail showing an example where the inclusion of two species (*i* and *j*) in an ecological background results in lower function (solid lines) than the additive expectation (dashed lines). This difference (*ϵ*_*ij*_) indicates a functional interaction between the two species. Δ*F*_*j*_ is the functional effect of species *j* on the ecological background. (**C**) Based on theoretical results from quantitative genetics, we hypothesized that the FEE slope of a species *i* may be explained by a sum of its effective interactions with every other species *j*. The effective interaction of species *i* with species *j* (denoted 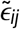) is defined as 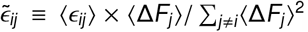, where the averages are taken across all possible ecological backgrounds where both species *i* and *j* are not present (*16,17*). (**D**) We quantified all effective interactions for every species and dataset (Materials and Methods), and used them to estimate the expected FEE slopes. The estimated slopes are in agreement with the empirical fits in Fig. 2 (*R*^2^ = 0.98). (**E**-**G**) We show three examples of species with positive, flat, and negative slopes. Slopes are explained by the sign and magnitude of the effective interactions between the focal species and its ecological partners.

To test whether this equation could explain the FEEs in Figs 1-2, we estimated the effective interactions between all pairs of species in those experiments (Materials and Methods). We found that eq. 1 does an excellent job at explaining the FEEs for all species across datasets (Fig. 3D, *R*^2^ = 0.98) and it helps us identify the mechanistic basis behind the different patterns of global epistasis for each species. For instance, *P. polymyxa* exhibits increasing returns, and this pattern is explained by its predominantly positive functional interactions with other community members (Fig. 3E). In turn, the flat slope exhibited by *Sphingoterrabacterium* sp. (Fig. 3F) arises from its evenly balanced positive and negative interactions, while the negative slope of *L. capitata* (Fig. 3G) can be attributed to its negative effective interactions with all other species.

The reader will have noticed that *P. polymyxa* displays two distinct types of functional effects on the function (amylolytic rate) of its background consortia, each described by a different FEE (Fig. 2D, rightmost panel). Closer examination of this species indicates that these two “branches” are defined by the presence or absence of a second species (*B. thuringiensis*) in the ecological background (fig. S4). The existence of these two branches can be rationalized from eq. 1 and the complementary equation that explains the intercept (fig. S5, Supplementary Text).

The analysis presented in Fig. 3 demonstrates that the FEE of a species can be explained mechanistically in terms of its ecological interactions, which in turn can be understood at the molecular level. For instance, as discussed above, the positive slope of *P. polymyxa* results from positive effective interactions with its ecological partners. These positive interactions have a known molecular basis: *P. polymyxa* is a biotin auxotroph whose growth is facilitated via cross-feeding by other members of the consortia (*25*). This observation highlights the utility of defining effective functional interactions between species in order to bridge the gap between molecular-level mechanisms and the emergence of community-level functions.

## FEEs can be leveraged to predict community-level functions

At the outset of this paper, we argued that the ability to predict the functional effects of a set of candidate species would allow us to quantitatively link the composition and function of any consortium one may form with them. In fact, FEEs make it possible to identify which of those consortia will optimize community function. A simple visual inspection of the FEEs can be useful for this purpose. For instance, species that exhibit patterns of *increasing costs* may be discarded due to their detrimental effect on community function, which is more pronounced in higher-performing backgrounds. On the other hand, species exhibiting *accelerating returns* could be more promising: they would act as functional “boosters” by bringing up the function of their background community, even more strongly when the background function is already high. Because FEEs are easily and intuitively interpretable, this simple observation can help us narrow down the list of potentially desirable species. Beyond these straightforward guidelines, we reasoned that FEEs could serve to obtain quantitative predictions of community function based on composition, and thus to identify optimal consortia.

Perhaps the simplest approach to predict the function of a consortium would be to iteratively concatenate the FEEs of all its members (Fig. 4A), as described in detail in the Materials and Methods and Supplementary Text. Despite its simplicity, this approach yields excellent results at predicting the function of newly assembled communities. For instance, in Fig. 4 we generated predictions for the community-level pyoverdine production of a set of 61 new consortia, none of which had been assembled in our previous experiment (i.e., in Fig. 1F). We then went back to the laboratory, assembled those consortia *de novo*, and measured their empirical function (fig. S1, Materials and Methods). The agreement between our predictions and the empirical measurements was excellent (*R*^2^ = 0.80, Fig. 4B). The success of this simple approach is also remarkable across all of the other datasets we had previously analyzed (fig. S6). The method was also able to successfully identify optimal consortia in our pyoverdines experiment (fig. S7) as well as all the others (fig. S8).

**Fig. 4.**
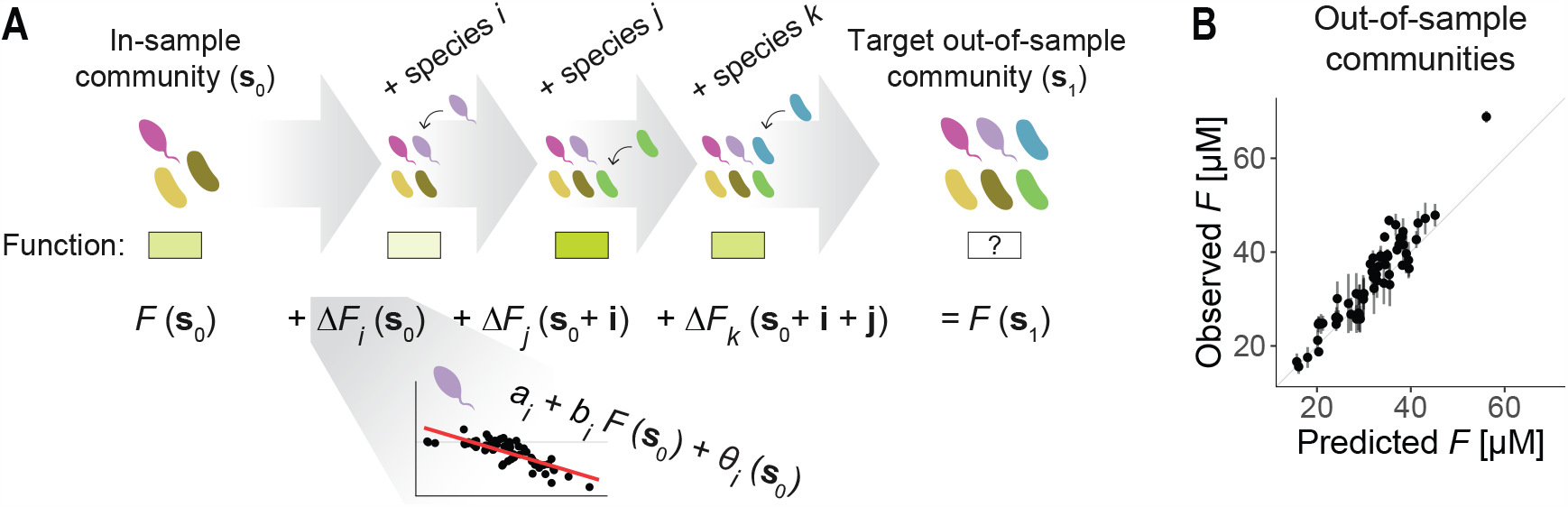
Ecological global epistasis can be leveraged to predict community functions. (**A**) We hypothesized that iteratively concatenating the functional effects of a set of species (here species *i, j* and *k*) could serve to predict the function of an out-of-sample community (**s**_1_) from that of an in-sample community (**s**_0_) (Materials and Methods). (**B**) We evaluated the performance of this method by assembling 61 new consortia (which served as the out-of-sample test set of communities) and comparing their predicted and measured levels of pyoverdines secretion. We found a good agreement (*R*^2^ = 0.80) between the observations and the predictions. Dots and error bars represent means and standard deviations across two biological replicates.

The observation that this very simple statistical approach can quantitatively predict community-level functions highlights the utility of characterizing the FEEs, as they provide a simple and interpretable pathway towards linking community composition and function. This also suggests that more sophisticated statistical machinery, recently developed in genetics to quantitatively predict phenotypes from genotypes (*18*–*22*), could be extended to ecology to predictively connect the composition and function of species assemblages (*31*).

## Discussion

Building predictive models of community function that integrate the full complexity of functional and ecological interactions is extremely challenging. Such models have only been built in a small number of case studies (*31*–*34*), but their parameterization required exhaustive empirical work that was highly specific to the taxa, environmental conditions, and functions being studied. Machine learning strategies are more scalable (*35,36*), but extracting interpretable biological information from them is generally difficult. As an alternative, a common coarse-grained description of ecological communities involves reducing community structure to a scalar metric of biodiversity (*37,38*). When averaged across multiple communities, biodiversity is often correlated with ecosystem function, but the variation is typically high and the specific form of this relationship can vary in different ecological contexts. The relationships between biodiversity and ecosystem function provide valuable insights in natural settings, but they cannot resolve which specific consortia one must form to maximize a function of interest.

Our findings suggest that these limitations can be overcome by leveraging an ecological analog to the concept of *global epistasis*, originally formulated in the context of genetics. We have shown that simple linear regression models (*functional effect equations* or FEEs) predict the functional effect of a species in a given background community, and they emerge ubiquitously across very different ecological conditions and collective functions. Much like genetic global epistasis patterns, FEEs can be characterized from a small subset of empirical observations (even in large combinatorial spaces) without the need for detailed information on the molecular mechanisms governing the interaction between every pair of species.

We propose that these FEEs may be interpreted as representations of emergent, coarse-grained species-by-community interactions. Historically, the study of ecological interactions has broken them down as the sum of pairwise species-by-species (s×s) and higher-order interactions (e.g., s×s×s, etc.) (*23*–*25,39*–*42*). This logic has paralleled the way in which genetic interactions have been traditionally partitioned, as the sum of pairwise gene-by-gene interactions (g×g), third order interactions (g×g×g), fourth order, and so on (*43*). The observation of global epistasis in genetic systems has revealed that epistasis can be instead partitioned into a *global* component, described by a linear regression between the fitness effect of a mutation and the fitness of the genetic background, and an idiosyncratic component described by the residuals of this fit. Building on recent analogies between genetic and functional ecological interactions (*23*–*25*), here we have demonstrated that the latter can be partitioned in the same manner, as the sum of a global species-by-community interaction (s×C) described by the FEEs, and an idiosyncratic component captured by the residuals. Furthermore, we have shown that the emergence of these global species-by-community interactions can be explained in terms of specific species-by-species interactions, expanding on recent theoretical results from the field of quantitative genetics (*16,17*).

Our results open up a plethora of new avenues for further investigation. For example, both in our own experiments and in the datasets we re-analyzed, community functions were quantified after a single batch. This type of scenarios are common in biotechnological applications: food fermentation or biofuel production typically occur in closed bioreactors over a predetermined time period, and crop fields are generally harvested once plants reach maturity. However, in other cases one might want to engineer ecological communities to be propagated in time, subject to environmental variation or the influx of invader species. It is unclear whether adding a new species to such communities (as opposed to simultaneously co-inoculating the new species together with the other members of its ecological background) may affect their ecological functions in a predictable manner. Another important consideration is the inoculum size. In our analyses, we have assessed the effects of including a given species in a community at a fixed density. It is reasonable to expect that the same species, inoculated at different starting population sizes, will have different effects on community function. In addition, here we have studied qualitatively simple community functions, which can be carried out to some degree by individual community members in isolation. Many applications may require the optimization of more complex functions (e.g., the simultaneous secretion of two or more metabolites, or the removal of multiple environmental pollutants). Clearly, additional work will be needed to shed light on these and other questions. This is likely to require generating combinatorial datasets mapping community structure to function with larger sets of species than the ones currently available. Yet, our observation that global epistasis-like patterns emerge in ecological communities has clear and immediate practical uses for community-level engineering. Perhaps most importantly, our results argue that predictively linking biological structure to function can be attained through a common, general quantitative formalism across scales of organization — from molecules and organisms to ecological communities.

## Acknowledgements

We thank M. Tikhonov, B. Ogbunugafor, M. Rebolleda-Gómez and all members of the Sanchez lab for helpful discussions. **Funding:** This work was supported by a Packard Foundation Fellowship to A.Sa., by the National Institutes of Health through grant 1R35 GM133467-01 to A.Sa., and by the Spanish Ministry of Science and Innovation through grant PID2021-125478NA-I00 to A.Sa. **Author contributions:** D.B. and A.Sa. conceived the study. J.D.-C. and A.Sk. analyzed data. J.D.-C. and A.Sa. designed experiments. J.D.-C. performed experiments. J.D.-C. and A.Sk. processed and analyzed experimental data. J.D.-C., A.Sk., J.C.C.V, D.B. and A.Sa. discussed and interpreted results. J.D.-C. and A.Sa. wrote the paper, with input from A.Sk, J.C.C.V and D.B. **Competing Interests:** The authors declare that no competing interests exist in relation to this manuscript. **Data and materials availability:** All data and code is available at https://github.com/jdiazc9/eco_global_epist.

## Supplementary Materials

Materials and Methods

Supplementary Text

Figs. S1 to S11

Tables S1 and S2

References (*44–48*)

## References

1. C. Wagg, S. F. Bender, F. Widmer, M. G. Van Der Heijden. Soil biodiversity and soil community composition determine ecosystem multifunctionality. Proceedings of the National Academy of Sciences 111(14), 5266–5270 (2014).

2. F. Lino, D. Bajic, J. C. Vila, A. Sanchez, M. Sommer. Complex yeast–bacteria interactions affect the yield of industrial ethanol fermentation. Nature Communications 12, 1498 (2021).

3. R. C. Wright, V.-P. Friman, M. C. Smith, M. A. Brockhurst. Functional diversity increases the efficacy of phage combinations. Microbiology 167(12), 001110 (2021).

4. S. E. Wuest, R. Peter, P. A. Niklaus. Ecological and evolutionary approaches to improving crop variety mixtures. Nature Ecology & Evolution 5(8), 1068–1077 (2021).

5. C. Bank. Epistasis and adaptation on fitness landscapes. Annual Review of Ecology, Evolution, and Systematics 53, 457–479 (2022).

6. R. MacLean, G. Perron, A. Gardner. Diminishing returns from beneficial mutations and pervasive epistasis shape the fitness landscape for rifampicin resistance in pseudomonas aeruginosa. Genetics 186(4), 1345–1354 (2010).

7. A. I. Khan, D. M. Dinh, D. Schneider, R. E. Lenski, T. F. Cooper. Negative epistasis between beneficial mutations in an evolving bacterial population. Science 332(6034), 1193–1196 (2011).

8. H.-H. Chou, H.-C. Chiu, N. F. Delaney, D. Segrè, C. J. Marx. Diminishing returns epistasis among beneficial mutations decelerates adaptation. Science 332(6034), 1190–1192 (2011).

9. L. Perfeito, A. Sousa, T. Bataillon, I. Gordo. Rates of fitness decline and rebound suggest pervasive epistasis. Evolution 68(1), 150–162 (2014).

10. S. Kryazhimskiy, D. P. Rice, E. R. Jerison, M. M. Desai. Global epistasis makes adaptation predictable despite sequence-level stochasticity. Science 344(6191), 1519–1522 (2014).

11. S. Schoustra, S. Hwang, J. Krug, J. A. G. de Visser. Diminishing-returns epistasis among random beneficial mutations in a multicellular fungus. Proceedings of the Royal Society B: Biological Sciences 283(1837), 20161376 (2016).

12. J. Otwinowski, D. M. McCandlish, J. B. Plotkin. Inferring the shape of global epistasis. Proceedings of the National Academy of Sciences 115(32), E7550–E7558 (2018).

13. M. S. Johnson, A. Martsul, S. Kryazhimskiy, M. M. Desai. Higher-fitness yeast genotypes are less robust to deleterious mutations. Science 366(6464), 490–493 (2019).

14. X. Wei, J. Zhang. Patterns and mechanisms of diminishing returns from beneficial mutations. Molecular Biology and Evolution 36(5), 1008–1021 (2019).

15. C. W. Bakerlee, A. N. Nguyen Ba, Y. Shulgina, J. I. Rojas Echenique, M. M. Desai. Idiosyncratic epistasis leads to global fitness–correlated trends. Science 376(6593), 630–635 (2022).

16. G. Reddy, M. M. Desai. Global epistasis emerges from a generic model of a complex trait. eLife 10, e64740 (2021).

17. J. Diaz-Colunga, A. Skwara, K. Gowda, R. Diaz-Uriarte, M. Tikhonov, D. Bajic, A. Sanchez. Global epistasis on fitness landscapes. arXiv (2022).

18. P. A. Romero, A. Krause, F. H. Arnold. Navigating the protein fitness landscape with gaussian processes. Proceedings of the National Academy of Sciences 110(3), E193–E201 (2013).

19. A. Tareen, M. Kooshkbaghi, A. Posfai, W. T. Ireland, D. M. McCandlish, J. B. Kinney. MAVE-NN: learning genotype-phenotype maps from multiplex assays of variant effect. Genome Biology 23, 98 (2022).

20. J. Otwinowski. Biophysical Inference of Epistasis and the Effects of Mutations on Protein Stability and Function. Molecular Biology and Evolution 35(10), 2345–2354 (2018). ISSN 0737-4038.

21. Z. R. Sailer, S. H. Shafik, R. L. Summers, A. Joule, A. Patterson-Robert, R. E. Martin, M. J. Harms. Inferring a complete genotype-phenotype map from a small number of measured phenotypes. PLoS Computational Biology 16(9), e1008243 (2020).

22. P. D. Tonner, A. Pressman, D. Ross. Interpretable modeling of genotype–phenotype landscapes with state-of-the-art predictive power. Proceedings of the National Academy of Sciences 119(26), e2114021119 (2022).

23. A. L. Gould, V. Zhang, L. Lamberti, E. W. Jones, B. Obadia, N. Korasidis, A. Gavryushkin, J. M. Carlson, N. Beerenwinkel, W. B. Ludington. Microbiome interactions shape host fitness. Proceedings of the National Academy of Sciences 115(51), E11951–E11960 (2018).

24. H. tEble, M. Joswig, L. Lamberti, W. B. Ludington. High dimensional geometry of fitness landscapes identifies master regulators of evolution and the microbiome. bioRxiv (2021).

25. A. Sanchez-Gorostiaga, D. Bajić, M. L. Osborne, J. F. Poyatos, A. Sanchez. High-order interactions distort the functional landscape of microbial consortia. PLoS Biology 17(12), e3000550 (2019).

26. D. L. Mould, N. J. Botelho, D. A. Hogan. Intraspecies signaling between common variants of pseudomonas aeruginosa increases production of quorum-sensing-controlled virulence factors. mBio 11(4), e01865–20 (2020).

27. A. Stintzi, K. Evans, J.-m. Meyer, K. Poole. Quorum-sensing and siderophore biosynthesis in pseudomonas aeruginosa: lasrllasi mutants exhibit reduced pyoverdine biosynthesis. FEMS Microbiology Letters 166(2), 341–345 (1998).

28. S. E. Kuebbing, A. T. Classen, N. J. Sanders, D. Simberloff. Above-and below-ground effects of plant diversity depend on species origin: an experimental test with multiple invaders. New Phytologist 208(3), 727–735 (2015).

29. G. Ghedini, D. J. Marshall, M. Loreau. Phytoplankton diversity affects biomass and energy production differently during community development. Functional Ecology 36(2), 446–457 (2022).

30. S. Langenheder, M. T. Bulling, M. Solan, J. I. Prosser. Bacterial biodiversity-ecosystem functioning relations are modified by environmental complexity. PLoS ONE 5(5), e10834 (2010).

31. R. L. Clark, B. M. Connors, D. M. Stevenson, S. E. Hromada, J. J. Hamilton, D. Amador-Noguez, O. S. Venturelli. Design of synthetic human gut microbiome assembly and butyrate production. Nature Communications 12(1), 1–16 (2021).

32. Y. Chen, C.-J. Lin, G. Jones, S. Fu, H. Zhan. Enhancing biodegradation of wastewater by microbial consortia with fractional factorial design. Journal of Hazardous Materials 171(1-3), 948–953 (2009).

33. A. Eng, E. Borenstein. Microbial community design: methods, applications, and opportunities. Current Opinion in Biotechnology 58, 117–128 (2019).

34. K. Gowda, D. Ping, M. Mani, S. Kuehn. Genomic structure predicts metabolite dynamics in microbial communities. Cell 185(3), 530–546.e25 (2022).

35. J. Thompson, R. Johansen, J. Dunbar, B. Munsky. Machine learning to predict microbial community functions: an analysis of dissolved organic carbon from litter decomposition. PLoS ONE 14(7), e0215502 (2019).

36. K. Qu, F. Guo, X. Liu, Y. Lin, Q. Zou. Application of machine learning in microbiology. Frontiers in Microbiology 10, 827 (2019).

37. G. F. Midgley. Biodiversity and ecosystem function. Science 335(6065), 174–175 (2012).

38. A. Shade. Diversity is the question, not the answer. The ISME Journal 11(1), 1–6 (2017).

39. X. Guo, J. Q. Boedicker. The contribution of high-order metabolic interactions to the global activity of a four-species microbial community. PLoS Computational Biology 12(9), e1005079 (2016).

40. X. Guo, J. Boedicker. High-order interactions between species strongly influence the activity of microbial communities. Biophysical Journal 110(3), 143a (2016).

41. H. Mickalide, S. Kuehn. Higher-order interaction between species inhibits bacterial invasion of a phototrophpredator microbial community. Cell Systems 9(6), 521–533 (2019).

42. E. Korkmazhan, A. R. Dunn. High-order correlations in species interactions lead to complex diversity-stability relationships for ecosystems. Physical Review E 105, 014406 (2022).

43. M. B. Taylor, I. M. Ehrenreich. Higher-order genetic interactions and their contribution to complex traits. Trends in Genetics 31(1), 34–40 (2015).

