## Supplementary Materials for "Global epistasis and the emergence of ecological function"

**This PDF file includes:**

Materials and Methods
Supplementary Text
Figs. S1 to S11
Tables S1 and S2
References (*44–48*)

### Materials and Methods

**Bacterial species isolation from environmental samples.** Bacterial isolates were obtained from the communities described in ref. (44). In short, environmental samples (soil, leaves...) were collected from different geographical locations and used to inoculate eight identical synthetic habitats containing 500 µL of minimal medium with citrate as the only supplied carbon source. Communities were stabilized by serial passaging for 7 cycles of 48 h growth and 1:125 dilution such that species unable to grow in these conditions were filtered out. Communities were then diluted to $10^{-6}$ and streaked out on chromogenic agar plates (CHROMagar Mastitis GN). Eight individual colonies were selected such that they were distinguishable from one another in their morphologies. Isolates were streaked out two more times to filter out potential contamination from different species. Isolates were identified at the genus level by Sanger sequencing their 16S rRNA gene (Table S2) — all 16S sequences were less than 97% similar.

**Combinatorial community assembly.** Experiments were done using minimal medium containing 4.65 g/L $Na_2HPO_4 \times 2H_2O$ (Sigma-Aldrich), 3 g/L $KH_2PO_4$ (Fisher Scientific), 1 g/L $NH_4Cl$ (Fisher Scientific), 0.5 g/L NaCl (Fisher Scientific) supplemented with 2.5 g/L sodium citrate (Sigma-Aldrich), 0.1 mM $CaCl_2$ (Sigma-Aldrich), 2 mM $MgSO_4$ (Fisher Scientific) and 1% trace mineral supplement (v/v; ATCC; stock contains 0.5 g/L Ethylenediaminetetraacetic acid [EDTA], 3 g/L $MgSO_4 \times 7H_2O$, 0.5 g/L $MnSO_4 \times H_2O$, 1 g/L NaCl, 0.1 g/L $FeSO_4 \times 7H_2O$, 0.1 g/L $Co[NO_3]_2 \times 6H_2O$, 0.1 g/L $CaCl_2$ [anhydrous], 0.1 g/L $ZnSO_4 \times 7H_2O$, 0.01 g/L $CuSO_4 \times 5H_2O$, 0.01 g/L $AlK[SO_4]_2$ [anhydrous], 0.01 g/L $H_3BO_3$, 0.01 g/L $Na_2MoO_4 \times 2H_2O$, 0.001 g/L $Na_2SeO_3$ [anhydrous], 0.01 g/L $Na_2WO_4 \times 2H_2O$ and 0.02 g/L $NiCl_2 \times 6H_2O$). Starter cultures were prepared by resuspending a single colony of each isolate into individual 50 mL conical tubes (Falcon) containing 20 mL of medium, and allowing them to grow for 48 h at 30°C. Cultures were then fully homogenized, and communities were assembled in 96-deep well U-bottom plates (Greiner Bio-One) filled with 500 µL of medium per well by inoculating 1 µL of each starter monoculture into the corresponding wells — further details can be found in the Supplementary Text. Communities were then incubated still at 30°C for 48 h.

**Quantification of pyoverdines concentration.** After incubation, cultures were fully homogenized and growth was tracked by measuring the optical density (OD) at 620 nm of 100 µL in an AccuSkan FC plate reader (Fisher Scientific). Cells were then pelleted by centrifuging the plates at 3000 rpm for 25 min. Supernatants were collected and filtered through multi-well 0.2 µm filters (Pall Corporation) by centrifuging at 3500 rpm for 5 min. The OD at 405 nm of 100 µL of the supernatants was quantified in the same plate reader. OD values were converted to units of concentration using a value of $1.9 \times 10^4$ $M^{-1}cm^{-1}$ for the extinction coefficient of pyoverdines at 405 nm (45).

**Quantification of functional effective interactions.** For each dataset we analyzed, the effective interaction between species $i$ and $j$ ($\tilde{\epsilon}_{ij}$) was quantified as explained in Fig. 3A-C and in the Supplementary Text, that is:

$$\tilde{\epsilon}_{ij} \equiv \langle \epsilon_{ij} \rangle \frac{\langle \Delta F_j \rangle_{B(i)}}{\sum_{j \neq i} \langle \Delta F_j \rangle^2_{B(i)}} \tag{S1}$$

The term $\epsilon_{ij}$ represents the deviation between the function of a community that contains both species $i$ and $j$ with respect to the additive expectation that they do not interact (Fig. 3B), that is, that the difference in function between communities $\mathbf{s}$ and $\mathbf{s}+\mathbf{i}+\mathbf{j}$ is the sum of the separate contributions of species $i$ and $j$. Mathematically, this means:

$$\epsilon_{ij} = \underbrace{F(\mathbf{s}+\mathbf{i}+\mathbf{j})}_{\substack{\text{function of}\\ \text{community containing}\\ \text{both } i \text{ and } j}} - \underbrace{\left[F(\mathbf{s}) + \Delta F_i(\mathbf{s}) + \Delta F_j(\mathbf{s})\right]}_{\substack{\text{additive}\\ \text{expectation}}} \tag{S2}$$

$$\epsilon_{ij} = F(\mathbf{s}+\mathbf{i}+\mathbf{j}) - \left[F(\mathbf{s}) + \underbrace{\left[F(\mathbf{s}+\mathbf{i}) - F(\mathbf{s})\right]}_{\Delta F_i(\mathbf{s})} + \underbrace{\left[F(\mathbf{s}+\mathbf{j}) - F(\mathbf{s})\right]}_{\Delta F_j(\mathbf{s})}\right] \tag{S3}$$

$$\epsilon_{ij} = F(\mathbf{s}+\mathbf{i}+\mathbf{j}) - F(\mathbf{s}+\mathbf{i}) - F(\mathbf{s}+\mathbf{j}) + F(\mathbf{s}) \tag{S4}$$

To quantify the average $\langle\epsilon_{ij}\rangle$ as it appears in eq. S1, we applied eq. S4 to every possible background community $\mathbf{s}$ not containing species $i$ nor $j$, and took the average over all $\epsilon_{ij}$. Note that not every dataset we analyzed is combinatorially complete, so in many instances some of the terms in eq. S4 might be unknown. In those cases, we computed the average over only the known values of $\epsilon_{ij}$.

The terms $\Delta F_j$ in eq. S1 represent the functional effect of species $j$ as defined in the main text, calculated simply as $F(\mathbf{s}+\mathbf{j}) - F(\mathbf{s})$. In eq. S1, these values appear averaged across those background communities that do not contain species $i$ (nor, naturally, species $j$) — this set of backgrounds is denoted as $B(i)$. Like before, whenever there were unknown terms due to incomplete data we averaged across the known values only.

**Iterative concatenation of FEEs to predict community functions.** We call $\mathbf{s}_0$ one of the empirically tested consortia (henceforth an *in-sample* community). One may then want to predict the function of an out-of-sample consortium (which we denote as $\mathbf{s}_1$). In the example shown in Fig. 4A, the out-of-sample consortium contains three more species ($i$, $j$ and $k$) than the in-sample one. Including the first species $i$ in the starting consortium $\mathbf{s}_0$ will have an effect in function that we can estimate from the linear FEE for species $i$: $\Delta F_i(\mathbf{s}_0) = a_i + b_i F(\mathbf{s}_0)$. The function of the consortium resulting from the addition of species $i$ to $\mathbf{s}_0$ would then simply be $F(\mathbf{s}_0+\mathbf{i}) = F(\mathbf{s}_0) + \Delta F_i(\mathbf{s}_0) = a_i + (1+b_i) F(\mathbf{s}_0)$. We can next estimate the functional effect of including species $j$ on the "updated" background consortium $\mathbf{s}_0+\mathbf{i}$, and finally the functional effect of species $k$ on $\mathbf{s}_0+\mathbf{i}+\mathbf{j}$ (Fig. 4A):

$$F(\mathbf{s}_1) = \underbrace{F(\mathbf{s}_0)}_{\substack{\text{starting in-sample}\\ \text{community function}}} + \underbrace{\Delta F_i(\mathbf{s}_0)}_{\substack{\text{functional effect of}\\ \text{species } i\\ \text{on community } \mathbf{s}_0}} + \underbrace{\Delta F_j(\mathbf{s}_0+\mathbf{i})}_{\substack{\text{functional effect of}\\ \text{species } j\\ \text{on community } \mathbf{s}_0+\mathbf{i}}} + \underbrace{\Delta F_k(\mathbf{s}_0+\mathbf{i}+\mathbf{j})}_{\substack{\text{functional effect of}\\ \text{species } k\\ \text{on community } \mathbf{s}_0+\mathbf{i}+\mathbf{j}}} \tag{S5}$$

This iterative procedure ultimately gives a prediction for the out-of-sample community function $F(\mathbf{s}_1)$. By estimating the residuals of the FEEs using maximum likelihood, we can refine predictions and ensure that the order of species addition (e.g., *i-j-k*, *j-i-k*, *k-i-j*...) does not affect the predicted value — this is further discussed in the Supplementary Text.

**Data analysis.** All analyses were performed using R version 4.1.2. (46)

### Supplementary Text

#### Combinatorial assembly of bacterial communities in 96-well plates

This protocol illustrates how the set of 164 communities described in the main text was assembled for the pyoverdines secretion experiment. The protocol requires six plates for the assembly of the consortia (henceforth the *starting plates*, labeled S0 to S5), and nine plates where the communities are inoculated and grown (henceforth the *experimental plates*, labeled E0 to E8). Prior to the assembly, each isolate was allowed to grow for 48 h in minimal M9 citrate medium. The starting plate S0 contains 300 µL of each starter culture in the wells indicated below (wells colored in gray contain 300 µL of fresh medium instead, wells colored in white are left empty):

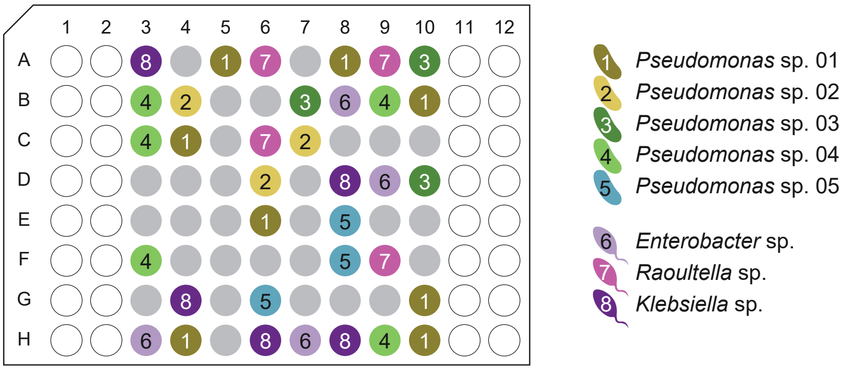


To assemble plates S1 to S5, we need to define a series of operations:

*Plate rotation*: transfer part of the volume from each well in a source plate (left in picture below) into a target plate (right in picture below), rotating well positioning 90 degrees clockwise:

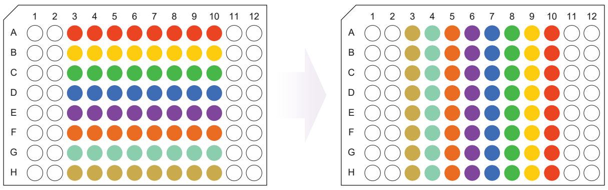

*Plate mixing*: transfer equal volumes from each well in a set of source plates (left in picture below) into a target plate (right in picture below):

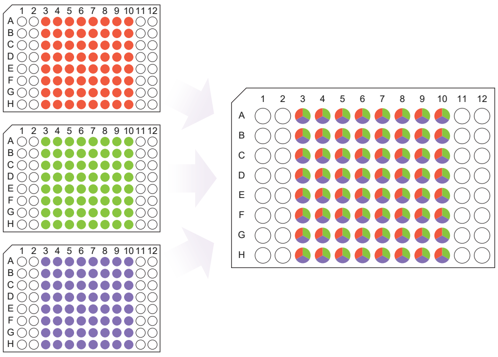

*Row re-arrangement*: transfer part of the volume from each well in a source plate (left in picture below) into a target plate (right in picture below), re-arranging well positions as indicated:

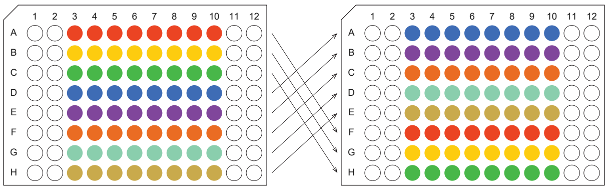

The starting plates S1 to S5 are assembled as follows:

plate S1 = *rotate* 200 µL per well from plate S0
plate S2 = *rotate* 100 µL per well from plate S1
plate S3 = *mix* plates S0, S1, and S2 (80 µL per well from each)
plate S4 = *re-arrange rows* of plate S3 (120 µL per well)
plate S5 = *mix* plates S3 and S4 (100 µL per well from each)

Plate S5 contains the inocula for our working set of background communities. The arrangement of these communities is as follows:

| | 1 | 2 | 3 | 4 | 5 | 6 | 7 | 8 | 9 | 10 | 11 | 12 |
|---|---|---|---|---|---|---|---|---|---|---|---|---|
| A | | | 1688 | 45 | 1458 | 1267 | 128 | 1478 | 1467 | 33678 | | |
| B | | | 11346 | 268 | 8 | 1 | 235 | 1256 | 2348 | 1 | | |
| C | | | 448 | 17 | 55 | 257 | 278 | 5 | 167 | 1144 | | |
| D | | | 148 | 485 | 567 | 1235 | 126 | 78 | 246 | 13477 | | |
| E | | | 13366 | 1167 | 18 | 18 | 2367 | 1258 | 134 | 138 | | |
| F | | | 14688 | 4 | 1458 | 2567 | 788 | 145 | 114677 | 13468 | | |
| G | | | 11144 | 2488 | 67 | 35 | 356 | 16 | 22448 | 1147 | | |
| H | | | 1346 | 11177 | 15 | 78 | 2367 | 18 | 14 | 11348 | | |

Where each number represents a different species: 1 - *Pseudomonas* sp. 01; 2 - *Pseudomonas* sp. 02; 3 - *Pseudomonas* sp. 03; 4 - *Pseudomonas* sp. 04; 5 - *Pseudomonas* sp. 05; 6 - *Enterobacter* sp.; 7 - *Raoultella* sp.; 8 - *Klebsiella* sp.

For the experiments, we filled 9 plates (E0 to E8) with 500 µL of minimal M9 citrate medium. We transferred 6 µL from plate S5 to each of these 9 plates (note that, thanks to the design of our protocol, this is equivalent to inoculating 1 µL of each corresponding monoculture). In addition, 1 µL of the monocultures of species 1 to 8 was inoculated into every well of plates E1 to E8 respectively — thus, plate E0 corresponds to the background communities, and plates E1 to E8 correspond to the addition of each species to those backgrounds. Communities where a same species was repeated multiple times (either in the background itself or after the addition of a knock-in) were disregarded for further analyses.

##### Maximum community functions across datasets

The combination of species that optimizes a particular function is not trivial to predict a priori based on the function of each taxon in isolation. For example, in our experiment, if there were no interactions we should expect that the consortium formed by the five pyoverdines-secreting species (*Pseudomonas* sp. 01 to 05) would maximize community function. Yet, we found that roughly 20% of the assembled consortia exhibited higher function than this community (fig. S1). In fact, the highest functional output in this experiment was achieved by a single species in monoculture (*Pseudomonas* sp. 01). However, it is worth noting that the functional maximum need not always correspond to a monoculture. In other experimental datasets, the best performing community contained multiple taxa (fig. S9), even including some whose function in monoculture was zero (e.g., *C. aerofaciens* in the Clark et al. dataset (*31*), Fig. 2E) or very small (e.g., *P. polymyxa* in the Sanchez-Gorostiaga et al. dataset (*25*), Fig. 2D).

##### Emergence of FEEs from functional interactions between species

As explained in the main text and the Materials and Methods, the functional effective interaction between two species $i$ and $j$ is denoted as $\tilde{\epsilon}_{ij}$ (eq. S1). By extending theoretical results from quantitative genetics (*16*,*17*) to the ecological scale, it follows that the FEE slope of a species $i$ would be given exactly by

$$b_i = \frac{\sum_{j\neq i} \langle \epsilon_{ij} \rangle \langle \Delta F_j \rangle_{B(i)} + \frac{1}{4} \sum_{k>j\neq i} \langle \epsilon_{ijk} \rangle \langle \epsilon_{jk} \rangle_{B(i)} + \cdots}{\sum_{j\neq i} \langle \Delta F_j \rangle^2_{B(i)} + \frac{1}{4} \sum_{k>j\neq i} \langle \epsilon_{jk} \rangle^2_{B(i)} + \cdots} \tag{S6}$$

Details on this expression can be found in refs. (*16*,*17*). Neglecting terms of order higher than two, and defining the effective interaction $\tilde{\epsilon}_{ij}$ as explained in the Materials and Methods (eq. S1), we arrive at eq. 1 in the main text, which estimates the FEE slope for species $i$ as $b_i \approx \sum_j \tilde{\epsilon}_{ij}$.

The FEE intercept ($a_i$) can be estimated as

$$a_i \approx \langle \Delta F_i \rangle_{B(i)} - b_i \langle F(\mathbf{s}) \rangle_{B(i)} \tag{S7}$$

where $\langle \Delta F_i \rangle_{B(i)}$ represents the average functional effect of species $i$ across all those background communities where $i$ is not present (we denote this set of backgrounds as $B(i)$). In turn, the term $\langle F(\mathbf{s}) \rangle_{B(i)}$ represents the average function of that same set of background communities where species $i$ is absent. Eq. S7 was used to quantify the expected intercepts of the FEEs shown in Fig. 3E-G (dashed lines). Note that eq. S7 is simply the general form of the least-squares linear regression intercept for a given slope $b_i$.

##### Branching in the FEEs for *P. polymyxa* can be explained by its effective functional interactions

Studying the effective interactions of *P. polymyxa* with its ecological partners can help us rationalize the "branching" observed in the FEE for this species (rightmost panel of Fig. 2D, fig. S4). As we mentioned in the main text, the two branches are determined by the presence or absence of *B. thuringiensis* in the ecological background (fig. S4). When we consider every background community (i.e., both those that contain and those that do not contain *B. thuringiensis*), we find that eqs. 1 and S7 give a pattern of diminishing returns, in very good agreement with the empirical regression when all backgrounds are accounted for (fig. S5, red linear fit). The negative slope is dominated by a strong negative functional interaction between *P. polymyxa* and *B. thuringiensis* (fig. S5).

Instead, we can consider only those background communities where *B. thuringiensis* is absent. In those backgrounds, the FEE for *P. polymyxa* is solely governed by its positive effective interactions with the other community members. We can use eqs. 1 and S7 to estimate the FEE slope and intercept for *P. polymyxa* in this subset of backgrounds: instead of averaging across all background communities, we can consider only those in which *B. thuringiensis* is absent. This gives rise to the rightmost branch of the FEE, with a positive slope that is in good agreement with the empirical fit (fig. S5, top panel). The remaining background communities all contain *B. thuringiensis.* Again, we can use eqs. 1 and S7, this time taking the averages only across those backgrounds in which *B. thuringiensis* is present. We find that the effective interactions of *P. polymyxa* with every other species remain positive, which explains the positive slope observed in this subset of backgrounds (fig. S5, bottom panel).

This second branch appears "shifted" leftwards with respect to the first one. This is because *B. thuringiesis* has, on average, a large positive functional effect in those background communities that do not contain *P. polymyxa*. Thus, the term $\langle F(\mathbf{s}) \rangle$ in eq. S7 is much larger when the average is taken across only those communities that contain *B. thuringiensis* than when it is taken across those communities where *B. thuringiensis* is absent. This makes the estimated intercept ($a_i$) substantially larger in the former case than in the latter, resulting in the apparent "split" (or "branching") between both sets of communities (fig. S5).

##### Estimation of FEE residuals to predict community function

As we discussed, we denote the intercept and slope of the FEE for species $i$ as $a_i$ and $b_i$ respectively. The functional effect of species $i$ on an ecological background $\mathbf{s}$ with function $F(\mathbf{s})$ is thus modeled as

$$\Delta F_i(\mathbf{s}) = a_i + b_i\, F(\mathbf{s}) + \theta_i(\mathbf{s}) \tag{S8}$$

where $\theta_i(\mathbf{s})$ represents the deviation from the fit (that is, the residual) corresponding to that background.

Our method for predicting community function is based on concatenating these FEEs as discussed in the main text (Fig. 4A) and the Materials and Methods. Suppose that we know the function of a specific *in-sample* community that we denote as $\mathbf{s}_0$, and we want to predict the function of the *out-of-sample* community resulting from the addition of one or various species to $\mathbf{s}_0$. We call this target community $\mathbf{s}_1$. In the example shown in Fig. 4A, $\mathbf{s}_1$ has three more species (that we generically denote as $i$, $j$ and $k$) than $\mathbf{s}_0$, but the method is applicable for any number of them. The function of $\mathbf{s}_1$ can be predicted by sequentially adding the functional effects induced by species $i$, $j$ and $k$ on the starting community $\mathbf{s}_0$:

$$F(\mathbf{s}_1) = \underbrace{F(\mathbf{s}_0)}_{\text{Starting in-sample community}} + \underbrace{\Delta F_i(\mathbf{s}_0)}_{\text{Addition of } i \text{ to } \mathbf{s}_0} + \underbrace{\Delta F_j(\mathbf{s}_0 + \mathbf{i})}_{\text{Addition of } j \text{ to } \mathbf{s}_0 + \mathbf{i}} + \underbrace{\Delta F_k(\mathbf{s}_0 + \mathbf{i} + \mathbf{j})}_{\text{Addition of } k \text{ to } \mathbf{s}_0 + \mathbf{i} + \mathbf{j}} \tag{S9}$$

where we have called $\mathbf{s}_0 + \mathbf{i}$ and $\mathbf{s}_0 + \mathbf{i} + \mathbf{j}$ to the intermediate communities resulting from adding species $i$, or both species $i$ and $j$, to the in-sample community $\mathbf{s}_0$. In order to quantify the values of the functional effects, we need the slopes and intercepts of the FEEs for species $i$, $j$ and $k$ (eq. S8). We can also estimate the values of the residuals $\theta_i(\mathbf{s}_0)$, $\theta_j(\mathbf{s}_0+\mathbf{i})$ and $\theta_k(\mathbf{s}_0+\mathbf{i}+\mathbf{j})$. Naturally, these residuals are unknown if the intermediate communities $\mathbf{s}_0+\mathbf{i}$ and $\mathbf{s}_0+\mathbf{i}+\mathbf{j}$ are not in-sample. Here we propose a method to estimate them using a maximum likelihood approach.

Note that the order in which species are added should not affect the final value of $F(\mathbf{s}_1)$, that is, the predicted function of the out-of-sample community should be independent from the order in which the functional effects are concatenated. Eq. S9 represents the case where the order of addition is $i \to j \to k$ (as illustrated in Fig. 4A), but any other order must yield the same value for $F(\mathbf{s}_1)$. This is because what we are here calling the "addition" of a species is not a dynamical process: For any species assemblage $\mathbf{s}$, we consider its function $F(\mathbf{s})$ to result from the simultaneous inoculation of all species in $\mathbf{s}$. Thus, the value of $F(\mathbf{s}_1)$ must match that in eq. S9 for any other order of "addition" (e.g., $j \to k \to i$). We call this the *closure condition* of the mapping between community structures and functions.

Let us focus on an arbitrary ecological background $\mathbf{s}$ and any pair of species $i$ and $j$. The closure condition tells us that the function of $\mathbf{s}+\mathbf{i}+\mathbf{j}$ should be the same regardless of which species' functional effect is considered first:

$$F(\mathbf{s}) + \Delta F_i(\mathbf{s}) + \Delta F_j(\mathbf{s}+\mathbf{i}) = F(\mathbf{s}) + \Delta F_j(\mathbf{s}) + \Delta F_i(\mathbf{s}+\mathbf{j}) \tag{S10}$$

Introducing the explicit form of the FEEs (eq. S8) into eq. S10 gives:

$$(1+b_i)\,\theta_j(\mathbf{s}) - (1+b_j)\,\theta_i(\mathbf{s}) + \theta_i(\mathbf{s}+\mathbf{j}) - \theta_j(\mathbf{s}+\mathbf{i}) = a_i b_j - a_j b_i \tag{S11}$$

which can be expressed in the following matrix form:

$$\begin{pmatrix} -(1+b_j) & (1+b_i) & 1 & -1 \end{pmatrix} \cdot \begin{pmatrix} \theta_i(\mathbf{s}) \\ \theta_j(\mathbf{s}) \\ \theta_i(\mathbf{s}+\mathbf{j}) \\ \theta_j(\mathbf{s}+\mathbf{i}) \end{pmatrix} = a_i b_j - a_j b_i \tag{S12}$$

This equation introduces constraints in the values of the residuals that ensure the closure condition is satisfied. Similar equations can be produced for any other background and pair of species — e.g., in our example with three species $i$, $j$ and $k$ and an in-sample background ($\mathbf{s}_0$) the constraints can be expressed in a compact form by defining matrix $\mathbf{M}$ and vectors $\mathbf{\Theta}$ and $\mathbf{C}$:

$$\underbrace{\begin{pmatrix} -(1+b_j) & (1+b_i) & 0 & -1 & 0 & 1 & 0 & 0 & 0 & 0 & 0 & 0 \\ 0 & -(1+b_k) & (1+b_j) & 0 & 0 & 0 & -1 & 0 & 1 & 0 & 0 & 0 \\ -(1+b_k) & 0 & (1+b_i) & 0 & -1 & 0 & 0 & 1 & 0 & 0 & 0 & 0 \\ 0 & 0 & 0 & 0 & 0 & 0 & 0 & -(1+b_j) & (1+b_i) & 1 & -1 & 0 \\ 0 & 0 & 0 & -(1+b_k) & (1+b_j) & 0 & 0 & 0 & 0 & 0 & 1 & -1 \\ 0 & 0 & 0 & 0 & 0 & -(1+b_k) & (1+b_i) & 0 & 0 & 1 & 0 & -1 \end{pmatrix}}_{\equiv \mathbf{M}} \cdot$$

$$\underbrace{\begin{pmatrix} \theta_i(\mathbf{s}_0) \\ \theta_j(\mathbf{s}_0) \\ \theta_k(\mathbf{s}_0) \\ \theta_j(\mathbf{s}_0+\mathbf{i}) \\ \theta_k(\mathbf{s}_0+\mathbf{i}) \\ \theta_i(\mathbf{s}_0+\mathbf{j}) \\ \theta_k(\mathbf{s}_0+\mathbf{j}) \\ \theta_i(\mathbf{s}_0+\mathbf{k}) \\ \theta_j(\mathbf{s}_0+\mathbf{k}) \\ \theta_i(\mathbf{s}_0+\mathbf{j}+\mathbf{k}) \\ \theta_j(\mathbf{s}_0+\mathbf{i}+\mathbf{k}) \\ \theta_k(\mathbf{s}_0+\mathbf{i}+\mathbf{j}) \end{pmatrix}}_{\equiv \mathbf{\Theta}} = \underbrace{\begin{pmatrix} a_i b_j - a_j b_i \\ a_j b_k - a_k b_j \\ a_i b_k - a_k b_i \\ a_i b_j - a_j b_i \\ a_j b_k - a_k b_j \\ a_i b_k - a_k b_i \end{pmatrix}}_{\equiv \mathbf{C}} \tag{S13}$$

Each row of $\mathbf{M}$ corresponds to a different pair of the three species ($i$-$j$, $j$-$k$ or $i$-$k$) in a different background ($\mathbf{s}_0$, $\mathbf{s}_0+\mathbf{i}$, $\mathbf{s}_0+\mathbf{j}$ or $\mathbf{s}_0+\mathbf{k}$). Any solution to eq. S13 will satisfy the closure condition. Analogous equations with the form $\mathbf{M} \cdot \mathbf{\Theta} = \mathbf{C}$ can be written for any number of species. This system of equations will generally be underdetermined (an infinite number of solutions will exist). Intuitively, not all of these are equally likely: solutions involving large values of the residuals would imply large deviations from the FEEs, and should in principle have low probability. We can formalize this intuition by studying the distribution of residuals of the fits for each linear FEE. For species $i$, this distribution will have a standard deviation $\sigma_i$ given by

$$\sigma_i^2 = \langle \theta_i^2 (\mathbf{s}) \rangle = \frac{1}{N} \sum_{\mathbf{s}} \left[ a_i + b_i\, F(\mathbf{s}) - \Big( F(\mathbf{s} + \mathbf{i}) - F(\mathbf{s}) \Big) \right]^2 \tag{S14}$$

where the sum goes over each of the $N$ possible ecological backgrounds $\mathbf{s}$ not containing species $i$. We can estimate $\sigma_i$ from partial observations if each FEE is fitted from a sufficiently large number of points, i.e., even if some terms of the sum are missing. We argue that the likelihood of a solution to eq. S13 will be dictated by the magnitude of the elements of $\mathbf{\Theta}$ in relation to the $\sigma_i$ corresponding to each of them. We define the estimated likelihood $\mathcal{L}$ of a vector $\tilde{\mathbf{\Theta}}$ that solves $\mathbf{M} \cdot \mathbf{\Theta} = \mathbf{C}$ (eq. S13) as

$$\mathcal{L}\left(\tilde{\Theta}\right) = \prod_{i,\mathrm{s}} \frac{1}{\sqrt{2\pi}\sigma_i} \exp\left( - \frac{\tilde{\theta}_i^2 (\mathbf{s})}{2\sigma_i^2} \right) \tag{S15}$$

where the $\tilde{\theta}_i (\mathbf{s})$ represent the elements of $\tilde{\mathbf{\Theta}}$ and the product goes over all species and ecological backgrounds. Note that eq. S15 would be exact only if the residuals were normally distributed and independent of one another. The first condition may not be met in general, and the second is not met since the closure condition imposes relationships among the residuals. However, eq. S15 can work as a useful approximation as it assigns low likelihood to solutions that involve values for the residuals that strongly deviate from the observed distributions.

It is convenient to define a scaled matrix $\mathbf{M}_\sigma$ and vector $\mathbf{\Theta}_\sigma$ as

$$\mathbf{\Theta}_\sigma = \mathbf{\Theta} \circ \begin{pmatrix} 1/\sigma_i \\ 1/\sigma_j \\ 1/\sigma_k \\ 1/\sigma_j \\ 1/\sigma_k \\ 1/\sigma_i \\ 1/\sigma_k \\ 1/\sigma_i \\ 1/\sigma_j \\ 1/\sigma_i \\ 1/\sigma_j \\ 1/\sigma_k \end{pmatrix} = \begin{pmatrix} \theta_i (\mathbf{s}_0) /\sigma_i \\ \theta_j (\mathbf{s}_0) /\sigma_j \\ \theta_k (\mathbf{s}_0) /\sigma_k \\ \theta_j (\mathbf{s}_0 + \mathbf{i}) /\sigma_j \\ \theta_k (\mathbf{s}_0 + \mathbf{i}) /\sigma_k \\ \theta_i (\mathbf{s}_0 + \mathbf{j}) /\sigma_i \\ \theta_k (\mathbf{s}_0 + \mathbf{j}) /\sigma_k \\ \theta_i (\mathbf{s}_0 + \mathbf{k}) /\sigma_i \\ \theta_j (\mathbf{s}_0 + \mathbf{k}) /\sigma_j \\ \theta_i (\mathbf{s}_0 + \mathbf{j} + \mathbf{k}) /\sigma_i \\ \theta_j (\mathbf{s}_0 + \mathbf{i} + \mathbf{k}) /\sigma_j \\ \theta_k (\mathbf{s}_0 + \mathbf{i} + \mathbf{j}) /\sigma_k \end{pmatrix} \tag{S16}$$

$$\mathbf{M}_\sigma = \mathbf{M} \circ \begin{pmatrix} \sigma_i & \sigma_j & \sigma_k & \sigma_j & \sigma_k & \sigma_i & \sigma_k & \sigma_i & \sigma_j & \sigma_i & \sigma_j & \sigma_k \\ \sigma_i & \sigma_j & \sigma_k & \sigma_j & \sigma_k & \sigma_i & \sigma_k & \sigma_i & \sigma_j & \sigma_i & \sigma_j & \sigma_k \\ \sigma_i & \sigma_j & \sigma_k & \sigma_j & \sigma_k & \sigma_i & \sigma_k & \sigma_i & \sigma_j & \sigma_i & \sigma_j & \sigma_k \\ \sigma_i & \sigma_j & \sigma_k & \sigma_j & \sigma_k & \sigma_i & \sigma_k & \sigma_i & \sigma_j & \sigma_i & \sigma_j & \sigma_k \\ \sigma_i & \sigma_j & \sigma_k & \sigma_j & \sigma_k & \sigma_i & \sigma_k & \sigma_i & \sigma_j & \sigma_i & \sigma_j & \sigma_k \\ \sigma_i & \sigma_j & \sigma_k & \sigma_j & \sigma_k & \sigma_i & \sigma_k & \sigma_i & \sigma_j & \sigma_i & \sigma_j & \sigma_k \end{pmatrix} \tag{S17}$$

where $\circ$ represents the Hadamard element-wise matrix product. Note that under this transformation, the product $\mathbf{M}_\sigma \cdot \mathbf{\Theta}_\sigma$ remains the same as $\mathbf{M} \cdot \mathbf{\Theta}$:

$$\mathbf{M}_\sigma \cdot \mathbf{\Theta}_\sigma = \mathbf{M} \cdot \mathbf{\Theta} = \mathbf{C} \tag{S18}$$

We can now calculate the Moore-Penrose pseudoinverse matrix of $\mathbf{M}_\sigma$, which we call $\mathbf{M}_\sigma^+$. The properties of this pseudoinverse make it so the following vector $\tilde{\mathbf{\Theta}}_\sigma^{(0)}$

$$\tilde{\mathbf{\Theta}}_\sigma^{(0)} = \mathbf{M}_\sigma^+ \cdot \mathbf{C} \tag{S19}$$

is the solution of the equation $\mathbf{M}_\sigma \cdot \mathbf{\Theta}_\sigma = \mathbf{C}$ that minimizes the sum of squares of its elements — and thus maximizes the estimated likelihood in eq. S15. Having obtained $\tilde{\mathbf{\Theta}}_\sigma^{(0)}$ from eq. S19, it is straightforward to undo the scaling introduced in eq. S16 and obtain estimates for the most likely values of the residuals.

Finally, these estimated residuals can be used to predict the function of the target out-of-sample community using eqs. S8 and S9. Note that at this point, the order in which species functional effects are considered does not affect the prediction, since the estimated residuals satisfy the closure condition. Furthermore, we can apply the closure condition to *every* possible path from and to *every* pair of communities $\mathbf{s}_0$ and $\mathbf{s}_1$. This has the advantage of making the closure condition globally satisfied. In this case, not every residual needs to be estimated — some are known from the in-sample observations. Without loss of generality, we can rearrange the elements of $\mathbf{\Theta}$ and the columns of $\mathbf{M}$ so that the first $n$ elements of $\mathbf{\Theta}$ correspond to residuals that are unknown (and thus they are treated as incognitas that we want to estimate), while the remaining are known from the in-sample observations:

$$\mathbf{\Theta} = \begin{pmatrix} \theta_1 \\ \theta_2 \\ \vdots \\ \theta_n \\ \theta_{n+1} \\ \theta_{n+2} \\ \theta_{n+3} \\ \vdots \end{pmatrix} \begin{matrix} \left.\vphantom{\begin{matrix}\theta_1\\\theta_2\\\vdots\\\theta_n\end{matrix}}\right\} \text{Unknown residuals} \\ \left.\vphantom{\begin{matrix}\theta_{n+1}\\\theta_{n+2}\\\theta_{n+3}\\\vdots\end{matrix}}\right\} \text{Known residuals} \end{matrix} \tag{S20}$$

It is then straightforward to rewrite $\mathbf{\Theta}$ as

$$\mathbf{\Theta} = \begin{pmatrix} \theta_1 \\ \theta_2 \\ \vdots \\ \theta_n \\ \theta_{n+1} \\ \theta_{n+2} \\ \theta_{n+3} \\ \vdots \end{pmatrix} = \underbrace{\begin{pmatrix} \theta_1 \\ \theta_2 \\ \vdots \\ \theta_n \\ 0 \\ 0 \\ 0 \\ \vdots \end{pmatrix}}_{\equiv \mathbf{\Theta}_1} + \underbrace{\begin{pmatrix} 0 \\ 0 \\ \vdots \\ 0 \\ \theta_{n+1} \\ \theta_{n+2} \\ \theta_{n+3} \\ \vdots \end{pmatrix}}_{\equiv \mathbf{\Theta}_2} \tag{S21}$$

This makes it so the equation to solve ($\mathbf{M} \cdot \mathbf{\Theta} = \mathbf{C}$, eq. S13) turns into

$$\mathbf{M} \cdot \mathbf{\Theta}_1 = \mathbf{C} - \mathbf{M} \cdot \mathbf{\Theta}_2 \tag{S22}$$

where the right-hand side is known. At this point, the dimension of the problem can be reduced since every element in $\mathbf{\Theta}_1$ after the $n$-th is zero. We can find the most likely values for the non-zero elements of $\mathbf{\Theta}_1$ (i.e., the unknown residuals) using the Moore-Penrose pseudoinverse like we discussed before.

For very large combinatorial datasets, considering all paths between all possible pairs of communities might result in the matrix $\mathbf{M}$ being extremely large. Numerically computing its pseudoinverse can thus be computationally out of reach. This is the case for the Clark et al. dataset described in the main text (*31*). For this dataset, we simplified our method by assuming that every unknown FEE residual was zero. To predict the function of a target community $\mathbf{s}_1$, we first identified the closest in-sample community (in terms of the number of species "additions" needed to reach $\mathbf{s}_1$). If two or more communities were equally close, we considered all of them. We then went through every path to $\mathbf{s}_1$ (i.e., every possible "order of addition" of species). Since assuming that all residuals are zero leads to the closure condition not being met, each path generally gives a different predicted value for $F(\mathbf{s}_1)$. We averaged all of them to obtain a final prediction. In some cases, considering every path from the closest in-sample community to the target community was computationally out of reach: if the two communities differ in the presence of $n$ species, there are $n!$ different paths from one to the other (and e.g., $10! > 3 \cdot 10^6$). In these cases, we randomly chose 1000 paths and averaged the predictions across them.

##### Evaluation of prediction accuracy and robustness

Despite its simplicity, our method for predicting community functions (Fig. 4A) outperformed linear regression models for the same purpose. A linear regression model estimates the function of a community $\mathbf{s}$ as

$$F(\mathbf{s}) = \sum_i \beta_i s_i + \sum_{j>i} \beta_{ij} s_i s_j + \sum_{k>j>i} \beta_{ijk} s_i s_j s_k + \cdots \tag{S23}$$

where $s_i = 0, 1$ represents the presence or absence of species $i$ in the community $\mathbf{s}$, and the coefficients ($\beta$) can be fit to a subset of empirical observations via the linear regression. This type of statistical model has been used to reconstruct fitness landscapes in genetics (e.g., (47)), and it can be seen as directly following from the way in which ecological and genetic interactions have been traditionally partitioned: into pairwise, third-order, and increasingly higher-order interactions (23–25,39–43). On the other hand, the method we described in this work (see Fig. 4A in main text, and also the Materials and Methods and Supplementary Text) is based on concatenating species' functional effects, and thus explicitly leverages the partitioning of ecological interactions into a "global" and an "idiosyncratic" component — captured by the FEE and its residuals, respectively. We therefore asked whether the latter method could yield more accurate predictions of community function than the former.

To answer this question, we compared the performance of our FEE-based method with that of first- and second-order regression models. We considered every dataset previously analyzed except for Clark et al. (31) (since, as explained above, inferring FEE residuals was not possible for this dataset). We then performed leave-one-out cross-validation by leaving each community out-of-sample one at a time, and using both our method and a linear regression method to predict its function. To prevent the regression models from overfitting to the limited empirical data they are trained with, we employed a least absolute shrinkage and selection operator (LASSO) regression with 10-fold cross-validation, a standard regularization method (48). If any method predicted a function lower than zero, we set the prediction to zero (note that none of the functions considered can have negative values). We then assessed the quality of the model predictions by quantifying the $R^2$ between the predicted and empirical functional values. For datasets with experimental replicates, we calculated the $R^2$ between the predicted value and the mean observed value of each experimental community. We found that our method yielded a higher quality of fit (as measured by $R^2$) than both the first- and second-order regressions in all of the datasets we considered (fig. S10).

To test how consistently our method could predict community function out-of-sample when reducing the number of communities used to fit FEEs, we first considered all the communities in our pyoverdine experiment. We randomly chose a subset of consortia to be left out-of-sample, and we used the remaining (in-sample) communities to fit a FEE for each species. We used our predictive statistical method (Materials and Methods, Fig. 4A) to predict the function of the out-of-sample communities, and then compared them to their empirical functions. We repeated this multiple times, each leaving a different (randomly chosen) subset of communities out-of-sample. We found that reducing the number of in-sample communities used to fit the FEEs affected the predictive power of our method only moderately. Even when FEEs were fit to a very small number of points, the predictions were still reliable ($R^2 \sim 0.5$ between predictions and observations with as few as $N \sim 4$ points per FEE, fig. S7). Similarly, in other datasets we also found that our method could provide accurate predictions and identify high-functioning consortia when reducing the number of in-sample communities (fig. S8).

The observation that FEEs can be leveraged to predict community function out-of-sample shows that these patterns capture meaningful information pertaining the mapping between community compositions and functions, and do not emerge as a trivial consequence of regression to the mean (see next section of Supplementary Text).

##### Empirical FEEs are not a consequence of regression to the mean

How generally can FEEs be expected to emerge? Can any arbitrary mapping between community composition and function lead to $\Delta F$-vs-$F$ correlations? Perhaps counter-intuitively, a negative FEE slope should be seen if the association between composition and function were random. In this scenario, the functions of any two consortia differing in the presence of a single taxon would be completely uncorrelated, and they can be seen as independent "draws" from a generic distribution of functions. If the first draw gives a large value for the function, the second is likely to give a smaller one and vice-versa due to simple regression to the mean. Thus, the subtraction of the two random functions (namely $F_2 - F_1$) would be likely to be positive if $F_1$ was small and negative if $F_1$ was large, leading to a negative correlation between $F_2 - F_1$ and $F_1$.

To test this intuition, we randomized the pairing between communities and functions in our pyoverdines dataset. Consistent with our reasoning, we found that the functional effects and the background functions exhibited a negative correlation in the randomized data (fig. S11). Interestingly, though, the FEEs we fit to our empirical data were significantly different to those in the randomized control (fig. S11). Negative slopes around $-1$ are generically observed when the association between community composition and function is random, but significantly different slopes commonly emerge in many real ecological contexts (e.g., Fig. 1F and Fig. 2). Despite the existence of negatively sloped $\Delta F$-vs-$F$ correlations, randomizing the association between composition and function should eliminate, or at the very least severely diminish, the ability of FEEs to predict community function out of sample. Application of our predictive method to the randomized dataset yielded unsurprisingly poor results (fig. S11). Together, these realizations suggest that the observed FEEs in empirical datasets across ecosystems and functions are not a trivial consequence of having a bounded set of functional values. This randomization control provides a benchmark against which we can determine whether the empirical FEEs capture ecologically meaningful information regarding the topography of ecological structure-function maps.

#### Supplementary Figures

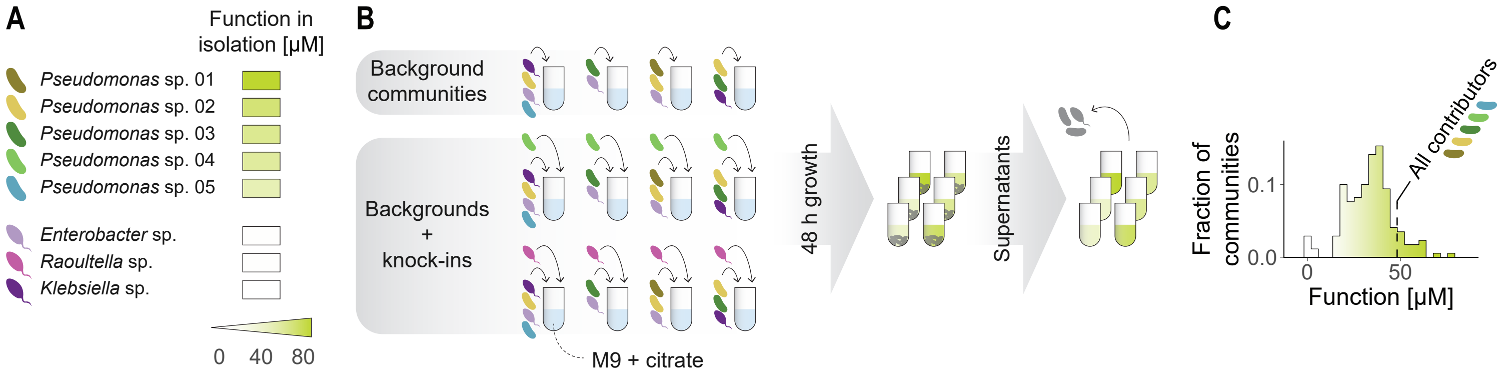


**Fig. S1. Assembly of microbial consortia in synthetic laboratory conditions.** (**A**) We isolated eight bacterial species from environmental samples and identified them at the genus level (Materials and Methods). Five of them exhibited secretion of pyoverdines when grown in monoculture in minimal M9 citrate medium (Materials and Methods). (**B**) We assembled 164 consortia by inoculating combinations of these eight species into minimal M9 citrate medium, and incubated them for 48 h. We then collected the spent media and quantified the concentration of pyoverdines in them (Materials and Methods). (**C**) We found variable levels of pyoverdines secretion, with the concentrations in the supernatants ranging from 0 to roughly 70 μM. About 20% of the assemblages exhibited higher function than the consortium formed by all five pyoverdines secretors.

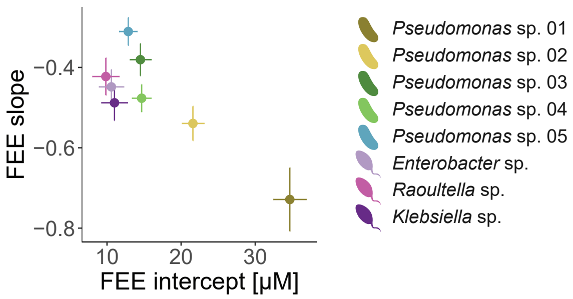


**Fig. S2. FEE slopes and intercepts vary across species.** Each species in our pyoverdine experiment exhibits a different FEE, characterized by its slope and intercept. Error bars represent standard deviations of the linear fit coefficients in Fig. 1F of the main text.

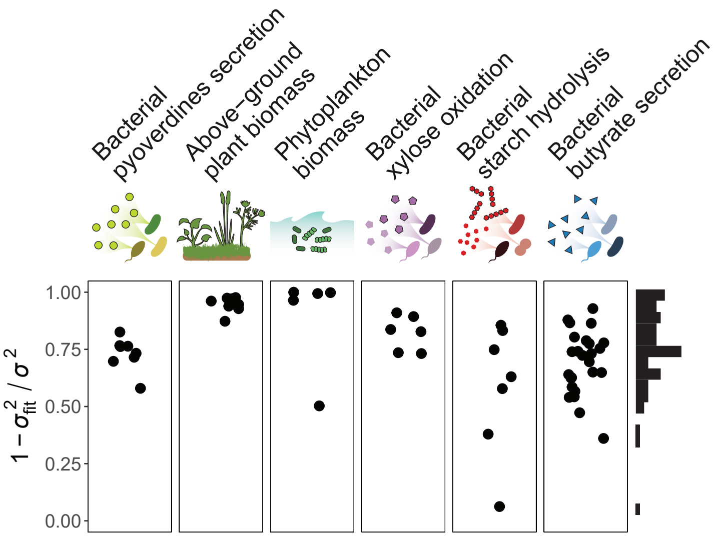


**Fig. S3. FEEs explain a high fraction of the variation in the functional effect of a species.** We fit a linear functional effect equation (FEE) to every species in each of the datasets described in the main text. The quality of the fits is here quantified as $1 - \sigma^2_{\text{fit}}/\sigma^2$, where $\sigma^2$ is the variance of the distribution of functional effects across all species in the dataset, and $\sigma^2_{\text{fit}}$ is the variance of the residuals of the fit for a given species. This is a metric of the degree to which the functional effect of a species is predictable from its FEE.

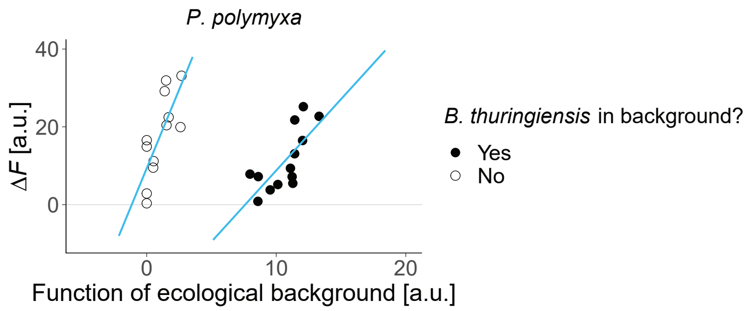


**Fig. S4. Branching in the functional effect equation of a species.** The effect on the amylolytic activity (*25*) of a community induced by *P. polymyxa* follows a different scaling with the function of the ecological background depending on whether *B. thuringiensis* is present (filled dots) or absent (hollow dots) in the background.

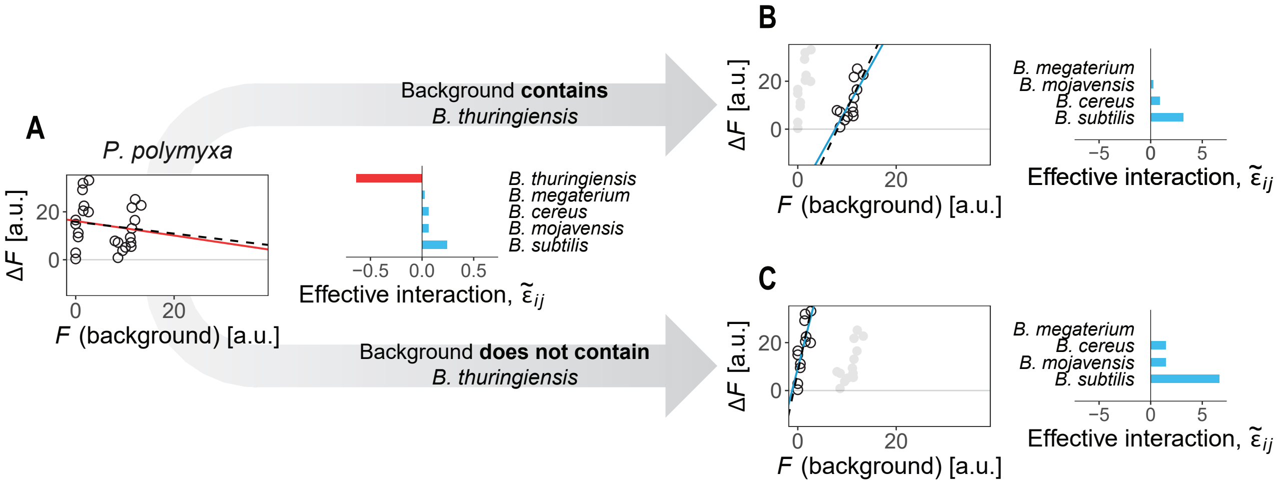


**Fig. S5. Effective interactions explain the branching observed in the FEEs for *P. polymyxa*.** (**A**) When we fit a single FEE for all backgrounds, we observe a negative slope. This can be explained by a strong negative effective interaction of *P. polymyxa* with *B. thuringiensis*. (**B**-**C**) When backgrounds are split by the presence/absence of *B. thuringiensis*, the effective interactions of *P. polymyxa* with the remaining species are positive, which gives rise to the positive FEE slopes in each branch. Dashed lines represent FEEs estimated using eqs. 1 and S7 (see main text and Supplementary Text), solid lines are empirical linear fits to the data.

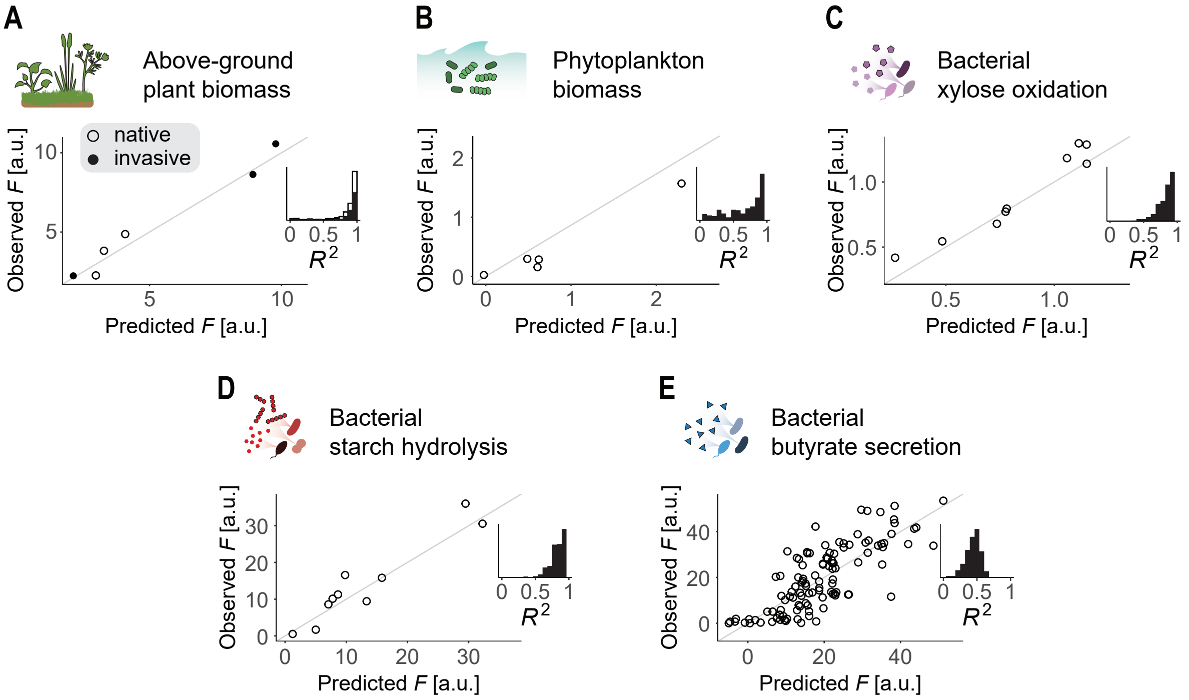


**Fig. S6. Predicting community function across datasets.** We evaluated the ability of our statistical method (Fig. 4A, Materials and Methods) to predict community functions in all datasets in table S1. For that, we left 20% of the communities in the datasets out of the sample, we used the remaining 80% to fit FEEs, and we applied our method to predict the function of the out-of-samples. We quantified the accuracy of the method as the $R^2$ between the predictions and the observations. We repeated the same process 500 times, each leaving a different subset of communities out of sample (randomly chosen). Main plots show an example of predicted against observed functions for one of the runs. Insets show histograms of the $R^2$ between predictions and observations across the 500 runs. (**A**) Data from Kuebbing et al. (*28*) Hollow/filled dots and bars correspond to native/invasive plants (note that this dataset is divided into two subsets of four plants each, see table S1). (**B**) Data from Ghedini et al. (*29*) (**C**) Data from Langenheder et al. (*30*) (**D**) Data from Sanchez-Gorostiaga et al. (*25*) (**E**) Data from Clark et al. (*31*)

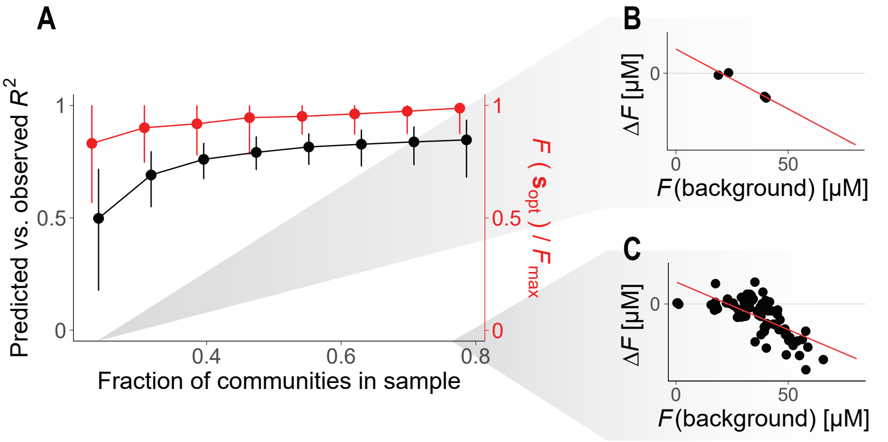


**Fig. S7. Predictions rely on the ability to accurately characterize FEEs.** We considered all the communities in our pyoverdine experiment. We left a subset of the communities out of the sample; the remaining (in-sample) communities were used to fit FEEs, and our predictive statistical method (Materials and Methods, Fig. 4A) was used to predict the function of the out-of-sample communities. The quality of the predictions was quantified as the $R^2$ between the predicted and observed functions of the out-of-sample communities (black line). We also evaluated the empirically measured function of the assemblage predicted to be optimal (here denoted as $\mathbf{s}_{opt}$), that is, to have the highest function out of those that we tested empirically (note that there are only 225 communities tested in our dataset — 164 in our first experiment and 61 additional ones in our second experiment — out of 255 total possible assemblages). We compared the measured function of this assemblage, $F(\mathbf{s}_{opt})$, with the true functional maximum, $F_{max}$ (red line). (**A**) The prediction method declined in accuracy as fewer communities were left in-sample, however, even for small sample sizes the signal remained strong and the predicted optimal community was either the true functional maximum or close to it. Dots represent means and error bars represent 95% confidence intervals across the 500 runs. (**B**) For the smallest sample sizes, the FEEs had to be estimated from a very small number of observations ($N \sim 4$ data points). (**C**) For larger sample sizes, FEEs were estimated from a high number of observations.

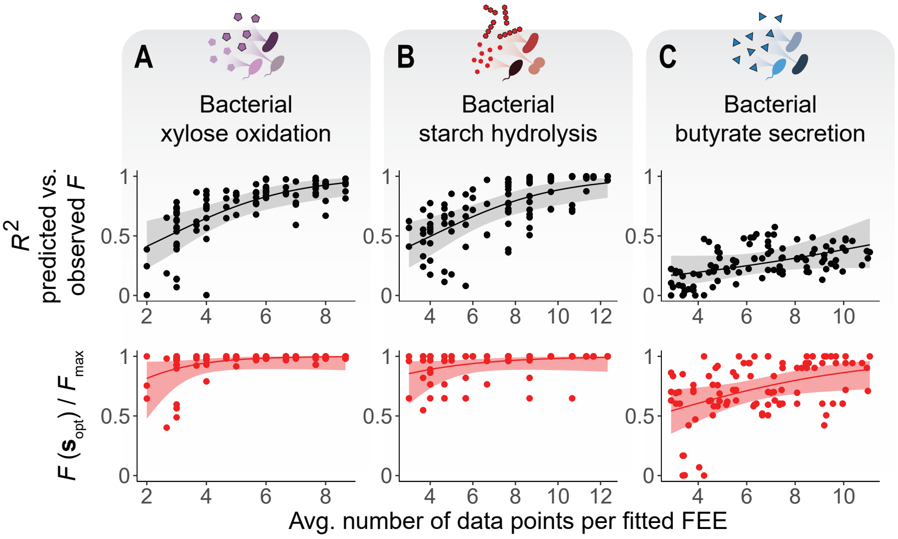


**Fig. S8. Prediction accuracy across datasets and sample sizes.** We repeated the analysis in fig. S7 for the three other datasets with the largest combinatorial size. We represent the number of data points used to fit each FEE, averaged across all species, against the $R^2$ between the predicted and measured functions of the out-of-sample communities (black), and against the function of the predicted functional maximum ($F(\mathbf{s}_{opt})$) with respect to the true maximum ($F_{max}$) across all measured assemblages (red). (**A**) Data from Langenheder et al. (*30*) (**B**) Data from Sanchez-Gorostiaga et al. (*25*) (**C**) Data from Clark et al. (*31*)

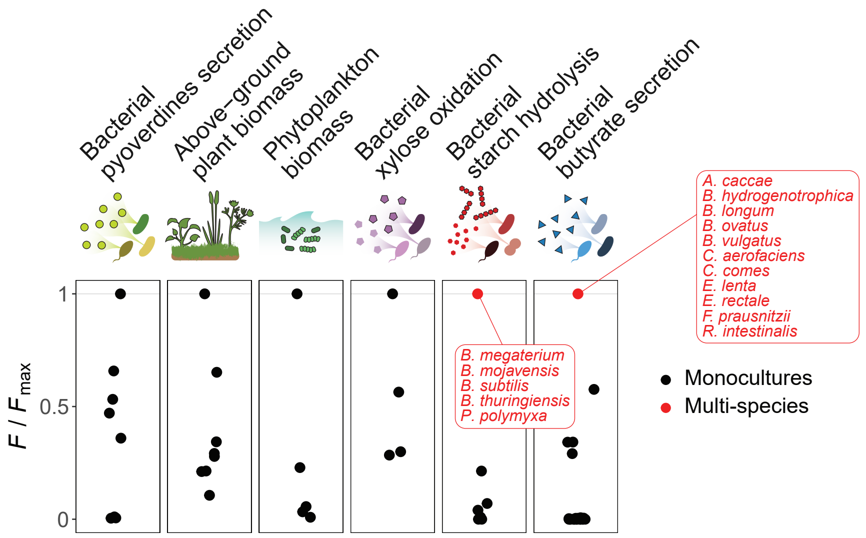


**Fig. S9. Optimal communities across datasets.** For each dataset, we represent the functions of the monocultures (black dots) with respect to the maximum function observed across all consortia ($F_{max}$). Whenever the functional maximum corresponds to a multi-species assemblage, it is also represented in red. Note that not all datasets are combinatorially complete, so the possibility that a multi-species community is the true functional maximum (which was not empirically tested) cannot be ruled out even if the one represented here is a monoculture.

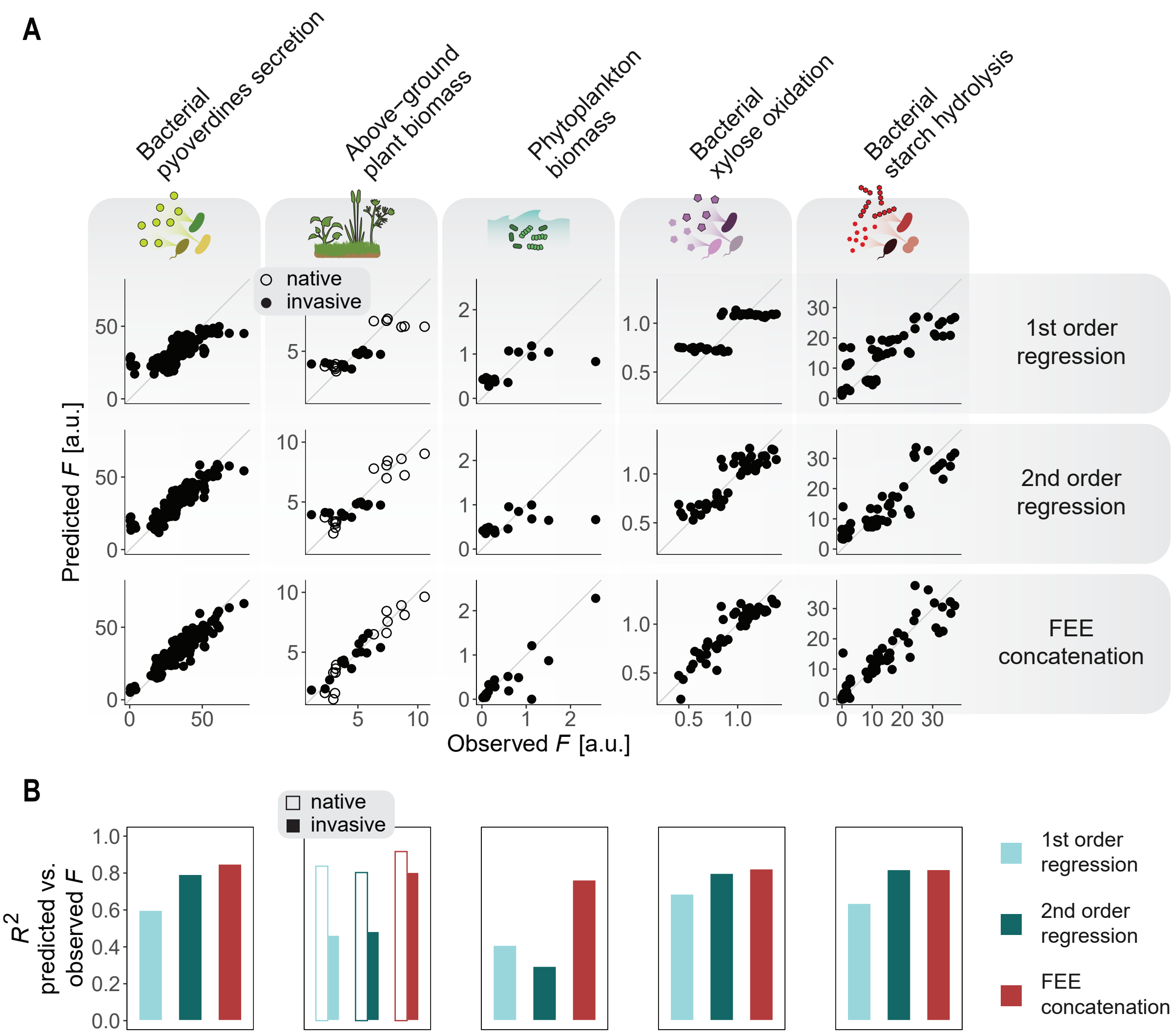


**Fig. S10. Predictions of community function based on FEEs concatenation are more accurate than alternative methods.** We performed leave-one-out cross-validations across various datasets to compare the performance of our predictive method with respect to first and second order regression models. (**A**) Each column of the grid corresponds to a different dataset, each row corresponds to a different method for predicting out-of-sample community functions. Note that in the Kuebbing et al. dataset (*28*) there are two subsets of species (a set of 4 native plants and another set of 4 invasive plants, see table S1), which were treated as separate datasets but are here represented together (with hollow/filled dots and bars, respectively). (**B**) The performance of each method was quantified as the $R^2$ between the predicted and observed values for the functions. Our method performed better, or as good as, than first or second order regressions.

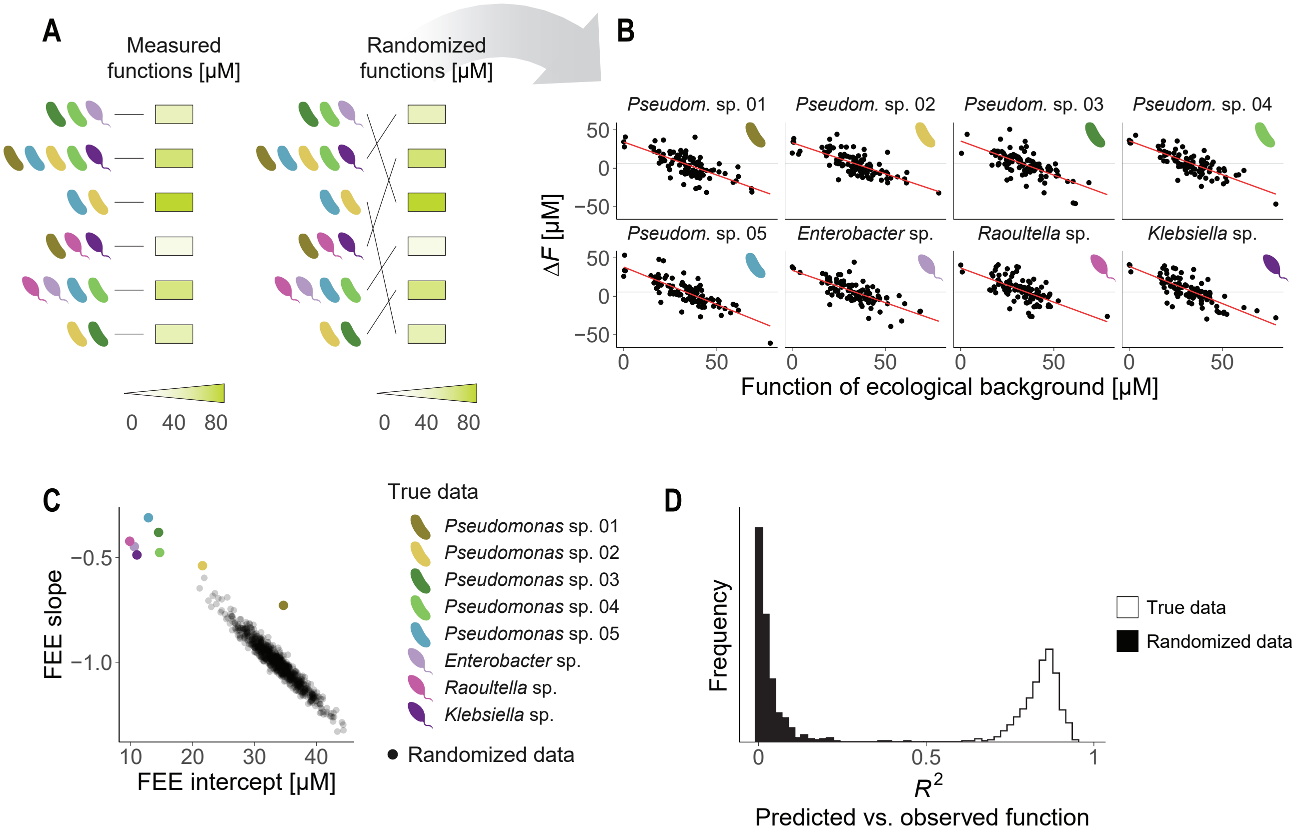


**Fig. S11. The observed FEEs indicate a non-random structure in the mapping between community compositions and functions.** (**A**) We randomized our data by rearranging the mapping between community structures and functions. (**B**) Correlations between the functions of the ecological backgrounds and the species' functional effects are observed in the randomized dataset. (**C**) We repeated the randomization process 500 times, each quantifying the slopes and intercepts of the FEEs (black dots). The true slopes and intercepts in the original data (colored dots) are not compatible with the ones found in the randomizations. (**D**) For each randomized set, we left 20% of the communities out of the sample and used the remaining 80% to fit FEEs and predict the out-of-sample functions using our statistical method (described in the Materials and Methods and Fig. 4A). This naturally yielded very poor results ($R^2 \sim 0$ between predictions and the observations). In contrast, the method had good performance ($R^2 \sim 0.8$) when applied to the true dataset.

#### Supplementary Tables

| Organisms type | Number of species | Ecosystem function | Source of dataset |
|---|---|---|---|
| Terrestrial plants | Two sets of 4 each | Above-ground biomass | Kuebbing et al. (28) |
| Phytoplankton | 5 | Biomass production | Ghedini et al. (29) |
| Bacteria | 6 | Xylose oxidation rate | Langenheder et al. (30) |
| Bacteria | 6 | Starch hydrolysis rate | Sanchez-Gorostiaga et al. (25) |
| Bacteria | 25 | Butyrate secretion | Clark et al. (31) |

**Table S1.** Summary of the datasets of combinatorial community composition and function analyzed in this study.

| Species | Sequence |
|---|---|
| *Pseudomonas* sp. 01 | CGCGCGTAGG TGGTTTGTTA AGTTGGATGT GAAAGCCCCG GGCTCAACCT<br>GGGAACTGCA TCCAAAACTG GCAAGCTAGA GTACGGTAGA GGGTGGTGGA<br>ATTTCCTGTG TAGCGGTGAA ATGCGTAGAT ATAGGAAGGA ACACCAGTGG<br>CGAAGGCGAC CACCTGGACT GATACTGACA CTGAGGTGCG AAAGCGTGGG<br>GAGCAAACAG GATTAGATAC CACGCCGTAA ACGATGTCAA CTAGCCGTTG<br>GAATCCTTGA GATTTTAGTG GCGCAGCTAA CGCATTAAGT TGACCGCCTG<br>GGGAGTACGG CCGCAAGGTT AAAACTCAAA TGAATTGACG GGGGCCCGCA<br>CAAGCGGTGG AGCATGTGGT TTAATTCGAA GCAACGCGAA GAACCTTACC<br>AGGCCTTGAC ATGCAGAGAA CTTTCCAGAG ATGGATTGGT GCCTTCGGGA<br>ACTCTGACAC AGGTGCTGCA TGGCTGTCGT CAGCTCGTGT CGTGAGATGT<br>TGGGTTAAGT CCCGTAACGA GCGCAACCCT TGTCCTTAGT TACCAGCACG<br>TAATGGTGGG CACTCTAAGG AGACTGCCGG TGACAAACCG GGGGGGATGA<br>CGTCAAGTCA TCATGGCCCT TACGGCCTGG GCTACACACG TGCTACAATG<br>GTCGGGTACA |
| *Pseudomonas* sp. 02 | CTGGGCGTAA GGTGGTTTGT TAAGTTGGAT GTGAAAGCCC CGGGCTCAAC<br>CTGGGAACTG CATCCAAAAC TGGCAAGCTA GAGTACGGTA GAGGGTGGTG<br>GAATTTCCTG TGTAGCGGTG AAATGCGTAG ATATAGGAAG GAACACCAGT<br>GGCGAAGGCG ACCACCTGGA CTGATACTGA CACTGAGGTG CGAAAGCGTG<br>GGGAGCAAAC AGGATTAGAT ACCCTGGTAG TCCACGCCGT AAACGATGTC<br>AACTAGCCGT TGGAATCCTT GAGATTTTAG TGGCGCAGCT AACGCATTAA<br>GTTGACCGCC TGGGGAGTAC GGCCGCAAGG TTAAAACTCA AATGAATTGA<br>CGGGGGCCCG CACAAGCGGT GGAGCATGTG GTTTAATTCG AAGCAACGCG<br>AAGAACCTTA CCAGGCCTTG ACATGCAGAG AACTTTCCAG AGATGGATTG<br>GTGCCTTCGG GAACTCTGAC ACAGGTGCTG CATGGCTGTC GTCAGCTCGT<br>GTCGTGAGAT GTTGGGTTAA GTCCCGTAAC GAGCGCAACC CTTGTCCTTA<br>GTTACCAGCA CGTTATGGTG GGCACTCTAA GGAGACTGCC GGTGACAAAC<br>CGGAGGAAGG TGGGGATGAC GTCAAGTCAT ACGGCCTGGG GCTACAATGG |
| *Pseudomonas* sp. 03 | GCGCGCGTAG GTGGTTTGTT AAGTTGGATG TGAAAGCCCC GGGCTCAACC<br>TGGGAACTGC ATCCAAAACT GGCAAGCTAG AGTACGGTAG AGGGTGGTGG<br>AATTTCCTGT GTAGCGGTGA AATGCGTAGA TATAGGAAGG AACACCAGTG<br>GCGAAGGCGA CCACCTGGAC TGATACTGAC ACTGAGGTGC GAAAGCGTGG<br>GGAGCAAACA GGATTAGATA CCCTGGTAGT CCACGCCGTA AACGATGTCA<br>ACTAGCCGTT GGAATCCTTG AGATTTTAGT GGCGCAGCTA ACGCATTAAG<br>TTGACCGCCT GGGGAGTACG GCCGCAAGGT TAAAACTCAA ATGAATTGAC<br>GGGGGCCCGC ACAAGCGGTG GAGCATGTGG TTTAATTCGA AGCAACGCGA<br>AGAACCTTAC CAGGCCTTGA CATGCAGAGA ACTTTCCAGA GATGGATTGG<br>TGCCTTCGGG AACTCTGACA TCAGCTCGTG TCGTGAGATG TTGGGTTAAG<br>TCCCGTAACG AGCGCAACCC TTGTCCTTAG TTACCAGCAC GTTATGGTGG<br>GCACTCTAAG GAGACTGCCG GTGACAAACC GGAGGAAGGG TGGGGGATGA<br>CGTCAAGTCA TCATGGCCCT TACGGCCTGG GCTACACACG TGCTACAATG |

| Species | Sequence |
|---|---|
| *Pseudomonas* sp. 04 | GCGCGTAGGT GGTTTGTTAA GTTGGATGTG AAAGCCCCGG GCTCAACCTG<br>GGAACTGCAT CCAAAACTGG CAAGCTAGAG TACGGTAGAG GGTGGTGGAA<br>TTTCCTGTGT AGCGGTGAAA TGCGTAGATA TAGGAAGGAA CACCAGTGGC<br>GAAGGCGACC ACCTGGACTG ATACTGACAC TGAGGTGCGA AAGCGTGGGG<br>AGCAAACAGG ATTAGATACC CTGGTAGTCC ACGCCGTAAA CGATGTCAAC<br>TAGCCGTTGG AATCCTTGAG ATTTTAGTGG CGCAGCTAAC GCATTAAGTT<br>GACCGCCTGG GGAGTACGGC CGCAAGGTTA AAACTCAAAT GAATTGACGG<br>GGGCCCGCAC AAGCGGTGGA GCATGTGGTT TAATTCGAAG CAACGCGAAG<br>AACCTTACCA GGCCTTGACA TGCAGAGAAC TTTCCAGAGA TGGATTGGTG<br>CCTTCGGGAA CTCTGACACA GGTGCTGCAT GGCTGTCGTC AGCTCGTGTC<br>GTGAGATGTT GGGTTAAGTC CCGTAACGAG CGCAACCCTT GTCCTTAGTT<br>ACCAGCACGT TATGGTGGGC ACTCTAAGGA GACTGCCGGT GACAAACCGG<br>AGGAAGGGTG GGGATGACG TCAAGTCATC ATGGCCCTTA CGGCCTGGGC<br>TACACACGTG CTACAATGGT |
| *Pseudomonas* sp. 05 | TGGGCGTAAA GCGCGCGTAG GTGGTTTGTT AAGTTGGATG TGAAATCCCC<br>GGGCTCAACC TGGGAACTGC ATCCAAAACT GGCAAGCTAG AGTATGGTAG<br>AGGGTGGTGG AATTTCCTGT GTAGCGGTGA AATGCGTAGA TATAGGAAGG<br>AACACCAGTG GCGAAGGCGA CCACCTGGAC TGATACTGAC ACTGAGGTGC<br>GAAAGCGTGG GGAGCAAACA GGATTAGATA CCCTGGTAGT CCACGCCGTA<br>AACGATGTCA ACTAGCCGTT GGGAGCCTTG AGCTCTTAGT GGCGCAGCTA<br>ACGCATTAAG TTGACCGCCT GGGGAGTACG GCCGCAAGGT TAAAACTCAA<br>ATGAATTGAC GGGGGCCCGC ACAAGCGGTG GAGCATGTGG TTTAATTCGA<br>AGCAACGCGA AGAACCTTAC CAGGCCTTGA CATCCAATGA ACTTTCCAGA<br>GATGGATTGG TGCCTTCGGG AGCATTGAGA CAGGTGCTGC ATGGCTGTCG<br>TCAGCTCGTG TCGTGAGATG TTGGGTTAAG TCCCGTAACG AGCGCAACCC<br>TTGTCCTTAG TTACCAGCAC GTAATGGTGG GCACTCTAAG GAGACTGCCG<br>GTGACAAACC GGAGGAAGGG TGGGGGATGA CGTCAAGTCA TCATGGCCCT<br>TACGGCCTGG GCTACACACG TGCTACAATG GTCGGTACAG |
| *Enterobacter* sp. | ACTGGGCGTA GGCGGTCTGT CAAGTCGGAT GTGAAATCCC CGGGCTCAAC<br>CTGGGAACTG CATTCGAAAC TGGCAGGCTA GAGTCTTGTA GAGGGGGGTA<br>GAATTCCAGG TGTAGCGGTG AAATGCGTAG AGATCTGGAG GAATACCGGT<br>GGCGAAGGCG GCCCCCTGGA CAAAGACTGA CGCTCAGGTG CGAAAGCGTG<br>GGGAGCAAAC AGGATTAGAT ACCCTGGTAG TCCACGCCGT AAACGATGTC<br>GACTTGGAGG TTGTGCCCTT GAGGCGTGGC TTCCGGAGCT AACGCGTTAA<br>GTCGACCGCC TGGGGAGTAC GGCCGCAAGG TTAAAACTCA AATGAATTGA<br>CGGGGGCCCG CACAAGCGGT GGAGCATGTG GTTTAATTCG ATGCAACGCG<br>AAGAACCTTA CCTACTCTTG ACATCCAGAG AACTTTCCAG AGATGGATTG<br>GTGCCTTCGG GAACTCTGAG ACAGGTGCTG CATGGCTGTC GTCAGCTCGT<br>GTTGTGAAAT GTTGGGTTAA GTCCCGCAAC GAGCGCAACC CTTATCCTTT<br>GTTGCCAGCG GTTCGGCCGG GAACTCAAAG GAGACTGCCA GTGATAAACT<br>GGAGGAGGGT GGGGGGATGA CGTCAAGTCA TCATGGCCCT TACGAGTAGG<br>GCTACACACG TGCTACAATG GCGCATACAA |

| Species | Sequence |
|---|---|
| *Raoultella* sp. | CTGGGCGTAA GCGCACGCAG GCGGTTTGTT AAGTCAGATG TGAAATCCCC<br>GGGCTCAACC TGGGAACTGC ATTTGAAACT GGCAAGCTTG AGTCTTGTAG<br>AGGGGGGTAG AATTCCAGGT GTAGCGGTGA AATGCGTAGA GATCTGGAGG<br>AATACCGGTG GCGAAGGCGG CCCCCTGGAC AAAGACTGAC GCTCAGGTGC<br>GAAAGCGTGG GGAGCAAACA GGATTAGATA CCCTGGTAGT CCACGCTGTA<br>AACGATGTCG ACTTGGAGGT TGTTCCCTTG AGGAGTGGCT TCCGGAGCTA<br>ACGCGTTAAG TCGACCGCCT GGGGAGTACG GCCGCAAGGT TAAAACTCAA<br>ATGAATTGAC GGGGGCCCGC ACAAGCGGTG GAGCATGTGG TTTAATTCGA<br>TGCAACGCGA AGAACCTTAC CTACTCTTGA CATCCAGAGA ACTTAGCAGA<br>GATGCTTTGG TGCCTTCGGG AACTCTGAGA CAGGTGCTGC ATGGCTGTCG<br>TCAGCTCGTG TTGTGAAATG TTGGGTTAAG TCCCGCAACG AGCGCAACCC<br>TTATCCTTTG TTGCCAGCGA TTCGGTCGGG AACTCAAAGG AGACTGCCAG<br>TGATAAACTG GAGGAAGGGG CATCATGGGC TAGGGCTACA |
| *Klebsiella* sp. | ATCGGATTAC CGCACGCAGG CGGTCTGTCA AGTCGGATGT GAAATCCCCG<br>GGCTCAACCT GGGAACTGCA TTCGAAACTG GCAGGCTGGA GTCTTGTAGA<br>GGGGGGTAGA ATTCCAGGTG TAGCGGTGAA ATGCGTAGAG ATCTGGAGGA<br>ATACCGGTGG CGAAGGCGGC CCCCTGGACA AAGACTGACG CTCAGGTGCG<br>AAAGCGTGGG GAGCAAACAG GATTAGATAC CCTGGTAGTC CACGCTGTAA<br>ACGATGTCGA CTTGGAGGTT GTTCCCTTGA GGAGTGGCTT CCGGAGCTAA<br>CGCGTTAAGT CGACCGCCTG GGGAGTACGG CCGCAAGGTT AAAACTCAAA<br>TGAATTGACG GGGGCCCGCA CAAGCGGTGG AGCATGTGGT TTAATTCGAT<br>GCAACGCGAA GAACCTTACC TACTCTTGAC ATCCACAGAA CTTAGCAGAG<br>ATGCTTTGGT GCCTTCGGGA ACTCTGAGAC AGGTGCTGCA TGGCTGTCGT<br>CAGCTCGTGT TGTGAAATGT TGGGTTAAGT CCCGCAACGA GCGCAACCCT<br>TATCCTTTGT TGCCAGCGGT CCGGCCGGGA ACTCAAAGGA GACTGCCAGT<br>GATAAACTGG GGGGATGACG TCAAGTCATC ATGGCCCTTA CGAGTAGGGG<br>CTACACACGT GCTACAATGG GCATATACAA |

**Table S2.** 16S rRNA gene sequences of the environmental isolates used in our experiment.

#### Supplementary References


44. J. Diaz-Colunga, N. Lu, A. Sanchez-Gorostiaga, C.-Y. Chang, H. S. Cai, J. E. Goldford, M. Tikhonov, Álvaro Sánchez. Top-down and bottom-up cohesiveness in microbial community coalescence. *Proceedings of the National Academy of Sciences* **119(6)**, e2111261119 (2022).

45. E. J. Drake, J. Cao, J. Qu, M. B. Shah, R. M. Straubinger, A. M. Gulick. The 1.8 å crystal structure of PA2412, an MbtH-like protein from the pyoverdine cluster of Pseudomonas aeruginosa. *Journal of Biological Chemistry* **282(28)**, 20425–20434 (2007).

46. R Core Team. *R: A Language and Environment for Statistical Computing*. R Foundation for Statistical Computing, Vienna, Austria (2022).

47. R. D. Kouyos, G. E. Leventhal, T. Hinkley, M. Haddad, J. M. Whitcomb, C. J. Petropoulos, S. Bonhoeffer. Exploring the complexity of the HIV-1 fitness landscape. *PLoS Genetics* **8(3)**, e1002551 (2012).

48. T. Hastie, R. Tibshirani, M. Wainwright. *Statistical Learning with Sparsity: The Lasso and Generalizations*. Chapman and Hall/CRC, 0 edition (2015). ISBN 978-0-429-17158-1.
